# Cell-free membrane systems reveal electrostatic control of ion transport in TMEM165 and disease-associated CaCA2 variants

**DOI:** 10.64898/2026.09.16.752219

**Authors:** Ekaterina Selivanovitch, Alexis R. Ostwalt, Angelica Byra, Zhongmou Chao, Alexandra Gaiffe, Guillemette van Raemdonck, Olivia Askowitz, Pierre Morsomme, Juliana Carten, Susan Daniel

**Author notes:** Equal contribution (Alexis R. Ostwalt, Angelica Byra).

## Abstract

Ion transport across intracellular membranes is essential for cellular homeostasis, yet organellar transport mechanisms remain difficult to resolve in native cells. Here, we establish a cell-free membrane system to interrogate full-length intracellular transporters under defined chemical conditions. Using cell-free protein synthesis, human Golgi transporter TMEM165 and its yeast homolog GDT1 were synthesized in membranes and analyzed by fluorescence and electrochemical measurements. The platform controlled protein identity, ion composition, and transmembrane pH gradients. TMEM165 transport was regulated by pH gradients, consistent with proton-coupled transport shaped by the electrostatic environment of the transport pathway. Disruption of a conserved acidic residue decreased pH sensitivity, broadened ion permeability, and altered steady-state transport behavior, indicating impaired proton coupling and selectivity. These findings identify conserved acidic residues as key regulators of TMEM165 function. More broadly, this work establishes a strategy for reconstructing organellar membrane proteins, resolving disease mechanisms, and engineering transport function in defined synthetic systems.

## Introduction

Transport proteins localized to intracellular compartments such as the Golgi, endosomes, and lysosomes operate within highly regulated and dynamic contexts, where ion gradients, trafficking pathways, and buffering systems are tightly coupled. As a result, functional readouts obtained in cells often reflect a convolution of direct transport activity with compensatory mechanisms, making it difficult to assign mechanistic roles to specific structural features and, from a translational perspective, study clinically relevant mutations. In this work, we address this challenge with two linked objectives: first, to establish and validate a cell-free membrane platform for assessing intracellular transporter function using the yeast transporter GDT1 and its human homolog TMEM165; and second, to use this platform to define how clinically associated TMEM165 mutations alter transport, pH sensitivity, and ion selectivity.

The study of membrane transporters has historically relied on systems in which proteins are accessible at the cell surface or can be readily purified and reconstituted without loss of function. However, even with currently employed reconstitution methods, transporters that reside within intracellular compartments remain poorly understood in part because of the difficulty of isolating their specific behavior from the surrounding cellular environment.^1, 2^ Cell-free protein synthesis (CFPS) provides a way to address this limitation by enabling intracellular transporters and clinically-relevant variants to be produced directly from DNA and inserted into defined environments without cellular expression, purification, or organelle isolation. This creates an opportunity to obtain rapid functional readouts of variant sequences while also resolving the physicochemical mechanisms that govern transport.

Examples of proteins that might benefit from the CFPS approach are members of the CaCA2 family, which represent a biologically important but mechanistically unresolved class of intracellular transporters. These conserved, multi-pass membrane proteins are proposed to function as cation-proton antiporters that maintain intracellular ion homeostasis by coupling divalent cation transport to proton gradients. Within this family, GDT1, a Saccharomyces cerevisiae protein localized to the Golgi and endomembrane system and has been functionally linked to Ca^2+^ and Mn^2+^ transport in model organisms.^3, 4^ Similarly, TMEM165, the human homolog of GDT1, is primarily localized to the Golgi, with additional localization reported in other intracellular compartments (e.g lysosomes), where it plays a critical role in regulating Mn^2+^, Ca^2+^, and proton homeostasis required for proper glycosylation.^5, 6^

Changes in TMEM165 function can lead to congenital disorders of glycosylation, highlighting the cell’s sensitivity to disrupted organelle ion homeostasis.^7-9^ At the molecular level, conserved acidic residues within the transmembrane region are thought to contribute to divalent cation coordination and transport, forming a key structural motif that is frequently altered in disease-associated variants.^10^ However, despite growing structural and computational insight, the transport mechanism of TMEM165 and the functional consequences of disease-associated mutations remain incompletely solved. A major challenge is the inability to isolate intrinsic transporter behavior from cellular context, where differences in the microenvironmental conditions across the membrane, lipid and protein composition within the membrane, and compensatory pathways can obscure direct functional effects. As a result, it remains difficult to determine whether a given disease-associated mutation disrupts transport chemistry itself, alters protein handling in cells, or produces a combination of both effects.

Deciphering the impact of specific mutations requires a controlled experimental framework in which wild-type (wt) and variant proteins can be compared directly under user-defined conditions. Variants of TMEM165, such as E108G, which directly disrupt the conserved acidic motif, are predicted to alter ion binding and electrostatic interactions, whereas others are thought to primarily affect localization or protein stability. In cellular systems, these effects are difficult to disentangle experimentally, as changes in trafficking or expression can obscure changes in underlying transport defects. Thus, resolving the TMEM165 ion transport mechanism, and its changes associated with mutations, requires not only a simplified membrane environment but also a method for producing and testing full-length transporters and its variants rapidly and under comparable conditions.

Simplified membrane systems alone provide a powerful approach for studying membrane transporters by reducing biological complexity and increasing its tailorability, while retaining the essential physical components required for transport. Synthetically-designed membrane platforms, including proteoliposomes and supported lipid bilayers (SLBs)^11, 12^, provide defined environments in which membrane proteins can be studied in isolation from the rest of the cell. In proteoliposomes, transport can be monitored by coupling ion flux to encapsulated reporters, allowing direct measurement of transmembrane movement under controlled gradients. ^13-15^ Supported lipid bilayers complement this approach by offering a planar geometry compatible with high-resolution imaging and electrical measurements, enabling analysis of insertion, topology, and transport-associated responses.^16-19^ Together these membrane mimetic systems establish a foundation for interrogating transport mechanisms outside the complexity of the cell.

However, membrane mimetics alone do not fully solve the problem of intracellular transporter access. Many *in vitro* membrane systems, particularly those formed from SLBs, rely on plasma membrane-derived vesicles, where proteins are accessible and can be harvested in their native context. In contrast, intracellular transporters localized to the Golgi and lysosomal compartments are significantly more difficult to isolate and reconstitute using these methods, as they are not directly accessible and often require extensive manipulation, including truncation or relocalization to the plasma membrane, to enable functional study. These limitations are especially important for TMEM and GDT1, where the goal is to compare full-length wild type and disease-associated variants under identical chemical conditions. Thus, an ideal platform would combine the chemical control of membrane mimetics with a protein-producing strategy that bypasses cellular expression and purification, such as CFPS.

CFPS provides a complementary strategy in which membrane proteins are produced directly from DNA templates in a reconstituted transcription-translation system and inserted into membrane bilayers without the need for purification or cellular expression.^20, 21^ Because new constructs can be generated directly from sequence information, CFPS is particularly well suited for rapid comparison of wild-type and patient-associated transporter variants. In recombinant cell-free systems, the transcriptional and translational machinery is supplied exogenously, enabling controlled protein production in the presence of liposomes or supported bilayers, where insertion occurs co-translationally or immediately following synthesis.^22, 23^ When coupled to proteoliposomes and SLBs, CFPS therefore extends minimal membrane systems from passive reconstitution platforms into programmable, sequence-defined systems for testing intracellular membrane proteins. This combined strategy is particularly advantageous for intracellular transporters, which are often difficult to express in living cells and may lose function when extracted using detergents. By coupling synthesis directly to membrane insertion, cell-free systems provide a route to preserve transporter integrity while enabling rapid and parallel testing of multiple constructs under identical conditions. Consistent with this, prior work has shown that cell-free supported membrane systems can preserve membrane protein activity as well. ^24, 25^

In this work, we first establish a bottom-up membrane platform that integrates cell-free expression with liposome and supported bilayer protein incorporation to directly interrogate ion transport by TMEM165 and GDT1. This approach enables rapid production and functional comparison of full-length intracellular transporters under defined chemical conditions (Fig. 1). We then use this platform to connect TMEM165 sequence variation to transporter behavior, with an emphasis on pH-dependent transport, clinically associated mutations, and ion selectivity. Using this approach, we find that TMEM165 activity is strongly dependent on transmembrane pH gradients, consistent with proton-coupled transport behavior governed by an electrostatic environment within the protein. We further characterize clinically associated TMEM165 variants linked to congenital disorders of glycosylation, including mutations that target the conserved acidic motif and others associated with altered transporter localization or stability. Notably, the disease-associated E108G mutation, which disrupts a key acidic residue within the proposed ion-coordination region, exhibits an increased apparent transport rate while simultaneously broadening ion permeability and altering steady-state ion transport. This behavior indicates that increased transport does not necessarily correspond to preserved function, but instead can reflect loss of coupling and ion selectivity. These results provide evidence that the electrostatic architecture of TMEM165 governs both substrate selectivity and responsiveness to chemical gradients. More broadly, comparison across variants reveals distinct differences in activity that would be difficult to disentangle in cellular contexts. By isolating transporter function under controlled conditions, this system enables variant-specific, molecular-scale analysis of how specific sequence features shape ion transport, selectivity, and coupling. Together, this work establishes a generalizable framework for dissecting intracellular membrane protein function and for connecting disease-associated mutations to their underlying physicochemical transport mechanisms.

**Figure 1.**
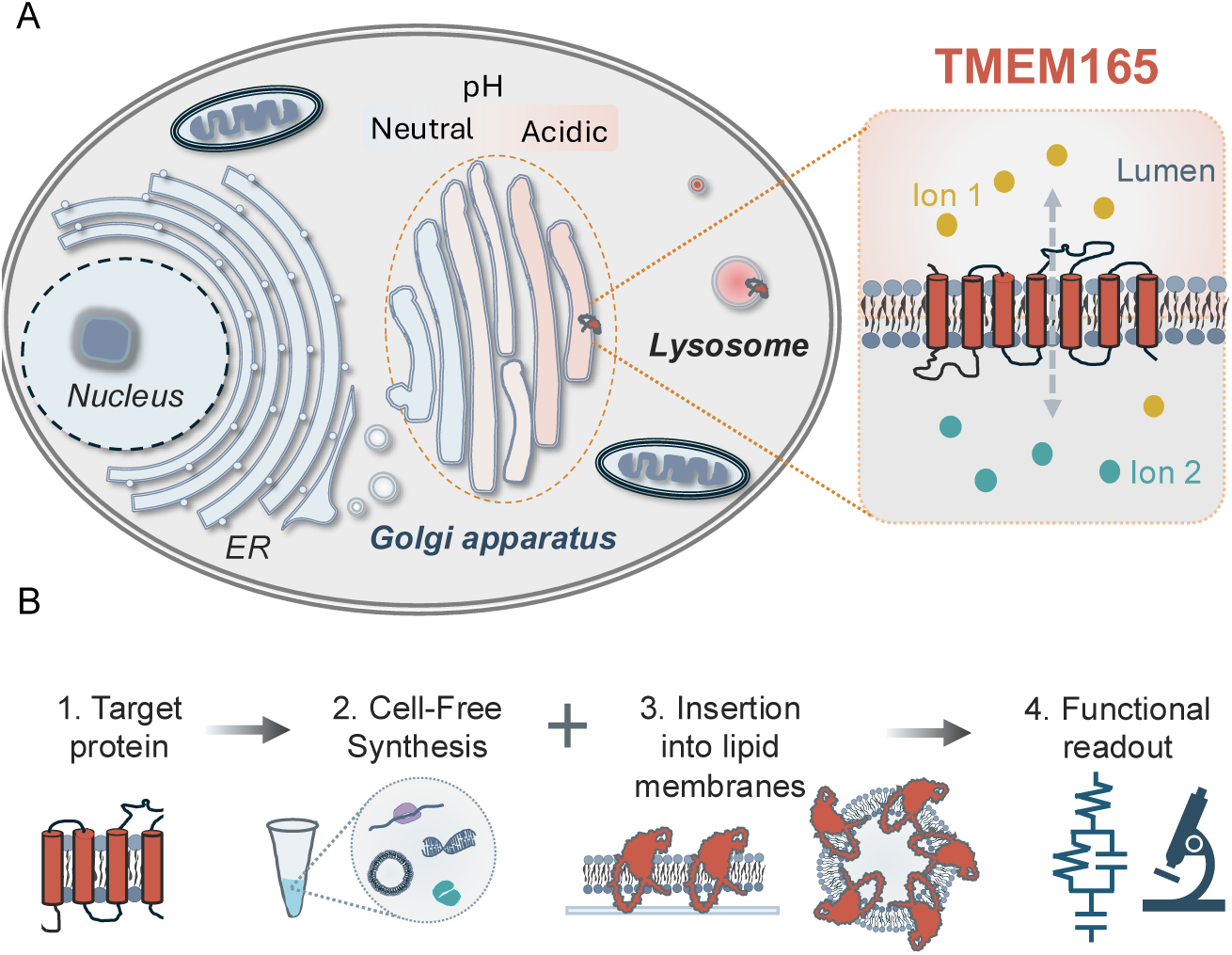
TMEM165 localization and cell-free membrane platform design. **(A)** TMEM165 localizes to the Golgi and lysosomal membranes, where it contributes to ion homeostasis and supports glycosylation. **(B)** Schematic overview of the platform used in this study, in which cell-free protein synthesis is used to synthesize and insert GDT1 and TMEM165 into supported lipid bilayers and liposomes. These platforms provide minimal, chemically defined systems for evaluating TMEM165 function.

## Results

### Platform Design

To establish a minimal and experimentally tractable platform for studying intracellular membrane transporters, we combined cell-free protein synthesis (CFPS) with complementary liposome- and supported lipid bilayer (SLB)-based membrane systems. Figure 1A provides cellular context for the class of organellar transporters examined in this study, highlighting their localization within intracellular membranes and their role in mediating ion and pH-dependent exchange across compartmentalized membrane barriers. Figure 1B outlines the general workflow used throughout this study, in which target transporters are synthesized by CFPS and inserted directly into synthetic lipid membranes for downstream functional interrogation.

For fluorescence-based measurements, liposomes containing encapsulated reporters sensitive to ion flux or pH were prepared prior to CFPS. Following synthesis in the presence of preformed liposomes, the resulting proteoliposomes provided a defined membrane environment in which intra- and extraliposomal conditions could be independently controlled. Size-exclusion chromatography separated reporter-loaded liposomes from free fluorophores, as verified by combined light-scattering and fluorescence measurements (Fig. S1 and S2). This configuration established a fluorescence-compatible system for imposing ion and pH gradients across the membrane.

To provide a complementary label-free membrane interface, we also developed an SLB-based platform integrated with conductive electrodes. Following bilayer formation, transporters were synthesized directly above the SLB, enabling insertion into a planar membrane architecture compatible with electrochemical impedance spectroscopy (EIS). Together, the liposome and SLB platforms provide orthogonal membrane formats for reconstituting intracellular transporters under defined experimental conditions and measuring their activity.

### GDT1 and TMEM165 as model CaCA2 transporters

We selected two CaCA2 transporters for this study: the yeast transporter GDT1 and its human homolog TMEM165. GDT1 provides a genetically and mechanistically defined reference system for establishing the cell-free membrane platform, whereas TMEM165 represents the disease-relevant human transporter. Both proteins contain conserved acidic sequence motifs characteristic of the CaCA2 family, including the E-φ-G-D-[KR]-[TS] motif positioned within membrane-spanning regions that are proposed to contribute to divalent cation coordination and proton-coupled transport (Fig. 2A, B). GDT1 contains six predicted transmembrane domains (TMDs) with a 23-amino acid N-terminal signal sequence. TMEM165 also contains six TMDs, with the conserved acidic motifs positioned within transmembrane helices 1 and 4. The TMEM165 signal domain is AA 1-33 of and it gets cleaved directly in the endoplasmic reticulum post insertion. For the purposes of this work, we maintained the full-length sequences of both constructs, including the signal peptide regions of both TMEM165 and GDT1. Together, these paired transporters allowed us to benchmark synthesis and membrane incorporation using a tractable yeast homolog before interrogating the human protein and clinically associated TMEM165 variants.

**Figure 2.**
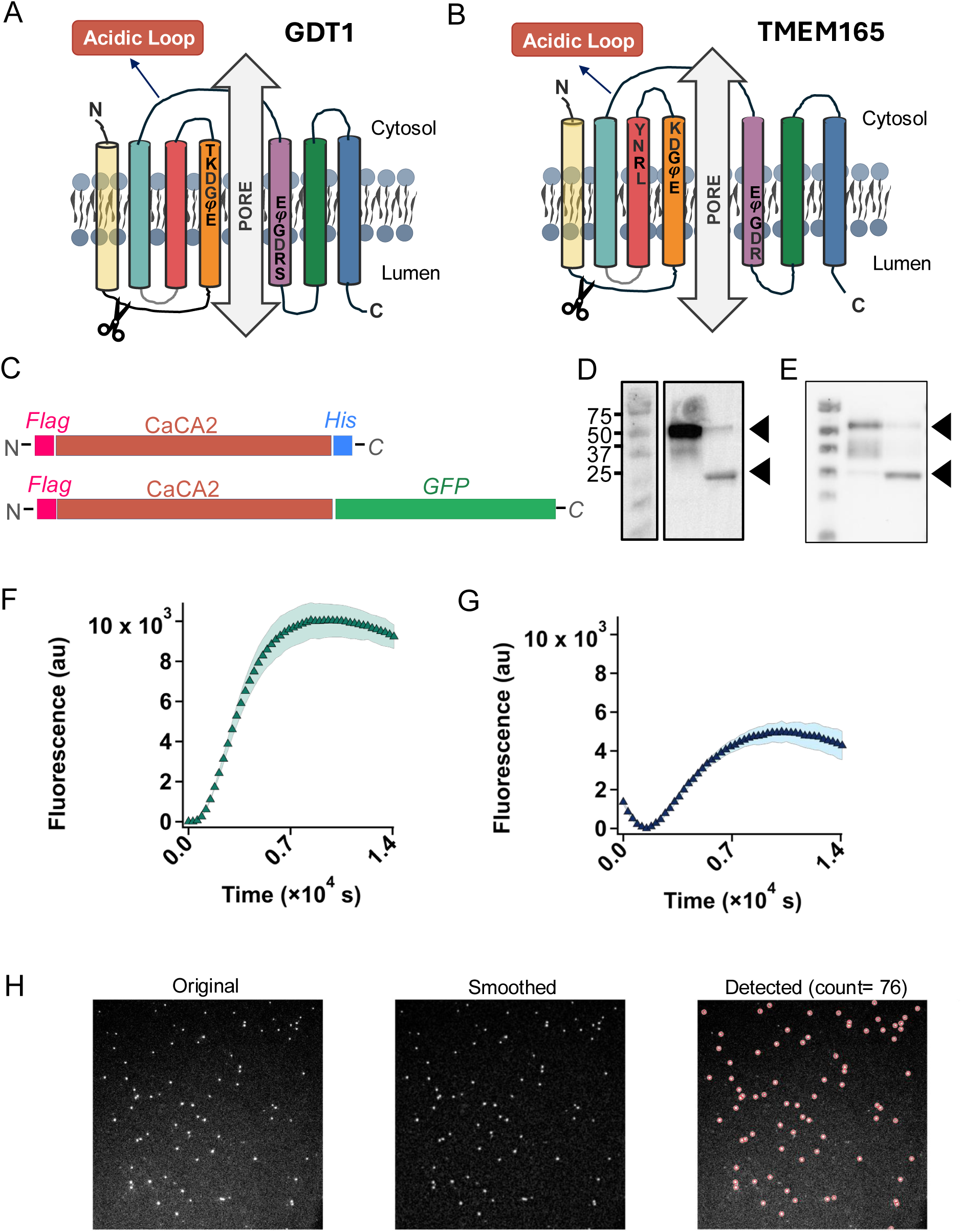
Cell-free synthesis and membrane association of GDT1 and TMEM165. **(A,B)** Schematic representations of the CaCA2 transporters (**A**) GDT1 and (**B**) TMEM165, showing predicted membrane topology, cytosolic and luminal orientation, conserved acidic-loop regions, and putative pore-forming regions. (**C**) Schematic of gene constructs used to assess cell-free protein synthesis, including epitope-tagged constructs and C-terminal GFP fusion constructs. (**D, E**) Anti-GFP western blot analysis of cell-free synthesized (**D**) GDT1-GFP and (**E**) TMEM165-GFP constructs. In each blot, the first lane shows the full-length transporter-GFP fusion construct, and the second lane shows the cleaved GFP-containing product after TEV addition. Arrowheads indicate the expected full-length and cleaved GFP species. Proteins were detected via Western blot using an anti-GFP primary antibody and an HRP-conjugated secondary antibody. (**F, G**) Time-dependent GFP fluorescence traces reporting cell-free synthesis of (**F**) GDT-GFP and (**G**) TMEM165-GFP. Data are shown as mean ± SD from n = 3 independent reactions. (**H**) TIRF microscopy analysis of TMEM165-GFP associated with supported lipid bilayers, showing the original fluorescence image, smoothed and enhanced image, and automated particle detection used to quantify SLB-associated fluorescent puncta.

### Cell-free synthesis and membrane association of CaCA2 transporters

GDT1 and TMEM165 were synthesized directly in the presence of preformed liposomes using cell-free protein synthesis. To monitor production, we generated fusion constructs in which GFP was appended to the C-terminus of either transporter (Fig. 2C). GFP fluorescence increased over time for both constructs, consistent with cell-free synthesis of the transporter fusion proteins (Fig. 2F, G). Protein production was further supported by western blot analysis, which showed bands at the expected molecular weights for the full-length fusion constructs, along with lower-molecular-weight species consistent with incomplete translation or partial degradation products (Fig. 2D, E).

We next asked whether the same cell-free strategy could be extended to planar membrane interfaces. GDT1-GFP and TMEM165-GFP were synthesized in the presence of supported lipid bilayers (SLBs), and membrane-associated fluorescence was evaluated using total internal reflection fluorescence (TIRF) microscopy. Because TIRF selectively excites fluorophores near the substrate-proximal membrane, the appearance of discrete fluorescent puncta within the evanescent field is consistent with local accumulation of transporter-GFP constructs at or near the SLB surface (Fig. 2H and Fig. S3, S4). Image processing and particle detection further enabled quantification of SLB-associated fluorescent species. Together, these results show that CFPS can produce full-length CaCA2 transporter constructs in both liposome and SLB membrane formats, establishing a foundation for subsequent functional measurements.

### Cell-free synthesis and activity in liposomes

Because the above constructs contain a GFP fusion, which may interfere with native folding and function of CaCA2 transporters, all functional studies were performed using constructs encoding only GDT1 or TMEM165 without GFP. For these constructs, protein expression was confirmed by western blotting, where bands corresponding to the expected molecular weights of ∼36–37 kDa for TMEM165 and ∼30–31 kDa for GDT1 were observed. Fractionation analysis further distinguished membrane-associated from non-associated protein populations (Figs. 3A and 3B), indicating that approximately 30–50% of the synthesized protein partitions into the membrane, consistent with typical yields for cell-free membrane protein expression systems.^26^

**Figure 3.**
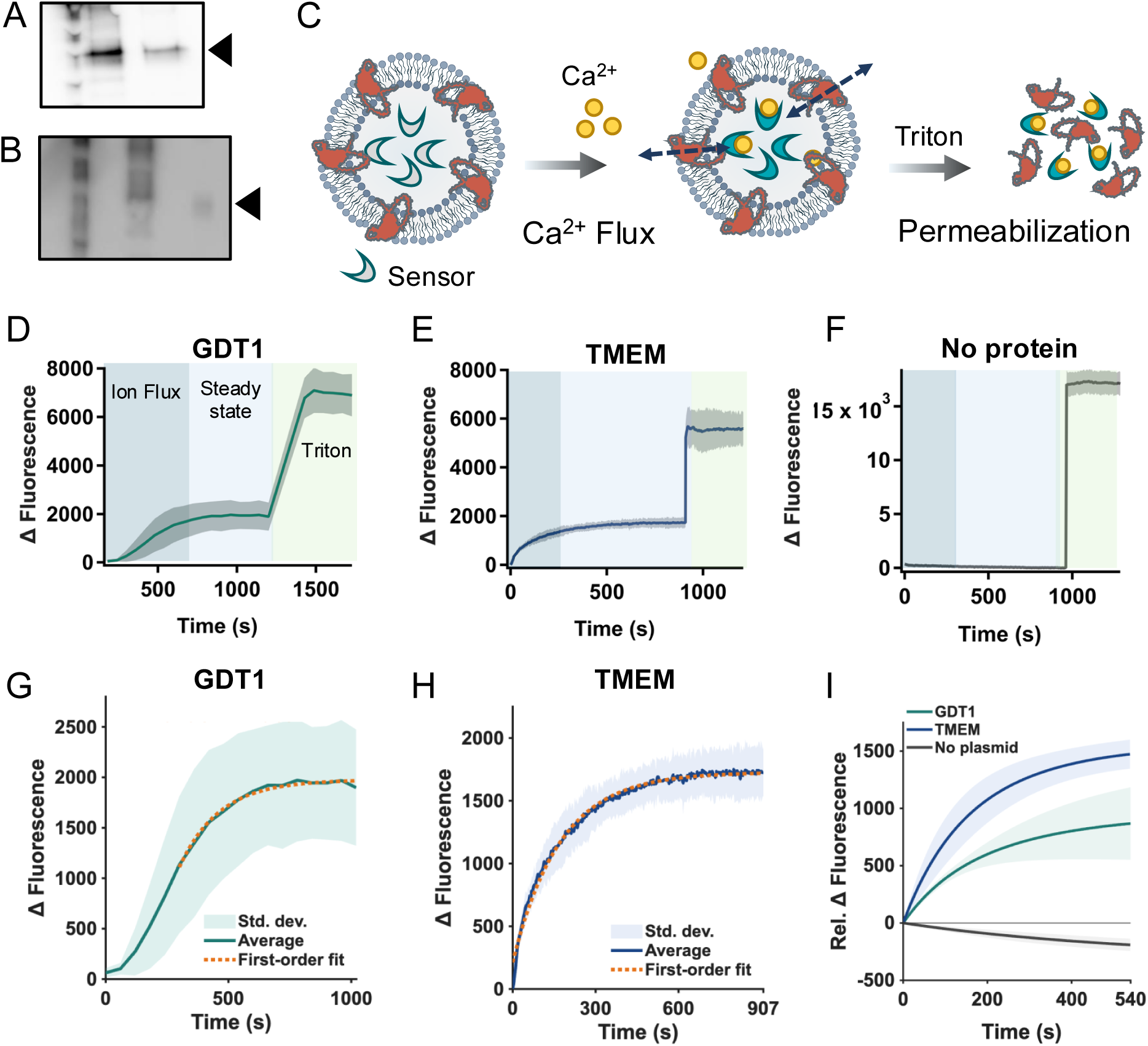
Cell-free proteoliposome assays enable functional characterization of GDT1- and TMEM165-mediated Ca²⁺ transport. (**A, B**) Western blot analysis of cell-free synthesized. Proteins were detected via Western blot using an anti-6x-His primary antibody and an HRP-conjugated secondary antibody (**A**) GDT1 and (**B**) TMEM165 before and after proteoliposome purification. In each blot, lane 1 shows the unpurified CFPS reaction and lane 2 shows the purified proteoliposome fraction. (**C**) Schematic of the liposome-based Ca²⁺ transport assay. GDT1 or TMEM165 was synthesized directly in the presence of liposomes containing encapsulated Fluo-4FF, a fluorescent Ca²⁺ reporter. Following external Ca²⁺ addition, Ca²⁺ influx into the liposomal lumen was detected as an increase in Fluo-4FF fluorescence. (**D–F**) Representative fluorescence traces from proteoliposomes containing (**D**) GDT1, (**E**) TMEM165, or (**F**) no protein following external Ca²⁺ addition. Protein-free liposomes showed minimal fluorescence change, supporting a protein-dependent transport response. (**G–I**) Initial transport phases from the corresponding traces were fit to a single-exponential function to compare Ca²⁺ transport kinetics. The time constant, τ, represents the characteristic time required for the fluorescence signal to approach its plateau; smaller τ values indicate faster Ca²⁺ influx. Data are shown for n = 3 independent replicates.

We next assessed transport activity using a fluorescence-based assay in proteoliposomes that contained wild-type GDT1 or TMEM165 (Fig. 3C). Ion influx was monitored through binding of Ca²⁺ to an encapsulated fluorescent sensor, enabling direct measurement of transport rates. To ensure quantitative interpretation, the linear dynamic range of the Fluo-4FF calcium sensor was first established (Fig. S5). Experimental conditions were selected such that Ca²⁺ was present in excess while the sensor remained below saturation, as sensor saturation would violate assumptions required for kinetic analysis.

For transport measurements, proteoliposomes containing either GDT1 or TMEM165 were introduced into buffer and Ca²⁺ was added externally (Fig. 3C). An initial rapid increase in fluorescence corresponds to Ca²⁺ binding to any free or externally accessible sensor, followed by a slower exponential increase attributed to Ca²⁺ transport across the membrane (Fig. 3D-E and Fig. S11-S13). Control liposomes lacking protein showed no measurable response (Fig. 3F), confirming that membrane transport is protein-dependent. To quantify this transport-associated phase, the fluorescence increase prior to detergent addition was fit using an apparent single-exponential rate equation (Fig. 3G, H). Because concentration-dependent transport measurements were not performed, this fit was used as an empirical description of the observed response rather than a full mechanistic kinetic model. The resulting apparent time constant, τ, describes the time required for the fluorescence signal to approach its asymptotic value under these assay conditions (suggesting all accessible encapsulated sensors bound to Ca^2+^), enabling comparison of GDT1- and TMEM165-mediated Ca²⁺ flux. This analysis assumes that Ca²⁺ binding to the sensor is rapid and effectively irreversible relative to membrane transport, such that the observed rate is dominated by ion translocation across the bilayer. Additional assumptions include negligible Ca²⁺ efflux, constant external Ca²⁺ concentration, and operation within the linear range of the sensor. Deviations from single-exponential behavior would indicate additional processes such as multiple transport populations, depletion of external Ca²⁺, or sensor saturation; however, under the conditions used here, a single-exponential model provides a consistent description of the data. Overlaying the fitted responses for GDT1, TMEM165, and protein-free liposomes further illustrates the protein-dependent nature of the transport response (Fig. 3I). To determine the maximum signal and estimate sensor occupancy, liposomes were permeabilized with Triton X-100, enabling complete equilibration of Ca²⁺ with the available sensors. This detergent-mediated maximum signal also allowed the transport-associated fluorescence increase to be expressed as a percentage of the total achievable fluorescence signal. Under these conditions, GDT1 and TMEM165 exhibited measurable activity, demonstrating that both are functionally active in this reconstituted system.

### Cell-free synthesis and activity in SLB

To assess the synthesis and membrane insertion of non-fluorescent GDT1 and TMEM165 constructs in SLBs we used immunostaining to detect exposed protein regions or epitope-specific tags. SLBs containing the respective cell-free synthesized proteins were incubated with fluorescent antibodies, and antibody binding was evaluated by total internal reflection fluorescence (TIRF) microscopy. The presence of fluorescent puncta indicates that the corresponding epitope is accessible at the bilayer-exposed surface and provides a readout of protein insertion and orientation.

For GDT1, the predicted topology places the acidic loop on the exposed side of the SLB, where it should be accessible to an anti-GDT1 antibody, whereas the C-terminal Strep tag is expected to face the opposite side of the bilayer and therefore be less accessible (Fig. 4A). However, because the shorter N-terminal hydrophobic sequence of GDT1 could potentially function as an additional membrane-spanning segment, the C-terminal Strep tag may instead be exposed on the accessible side of the SLB. Consistent with the predicted topology, anti-GDT1 staining produced a stronger punctate signal than anti-Strep staining (Fig. 4C,D). Detection of the Strep epitope may therefore reflect a minor population of oppositely oriented or incompletely inserted proteins, partial accessibility of the C terminus, or an alternative topology in which the C-terminal Strep tag is exposed. However, because the corresponding gel analysis showed few lower-molecular-weight products, the stronger anti-GDT1 signal is most consistent with a major population of GDT1 showing greater accessibility of the acidic loop than the C-terminal Strep tag.

**Figure 4.**
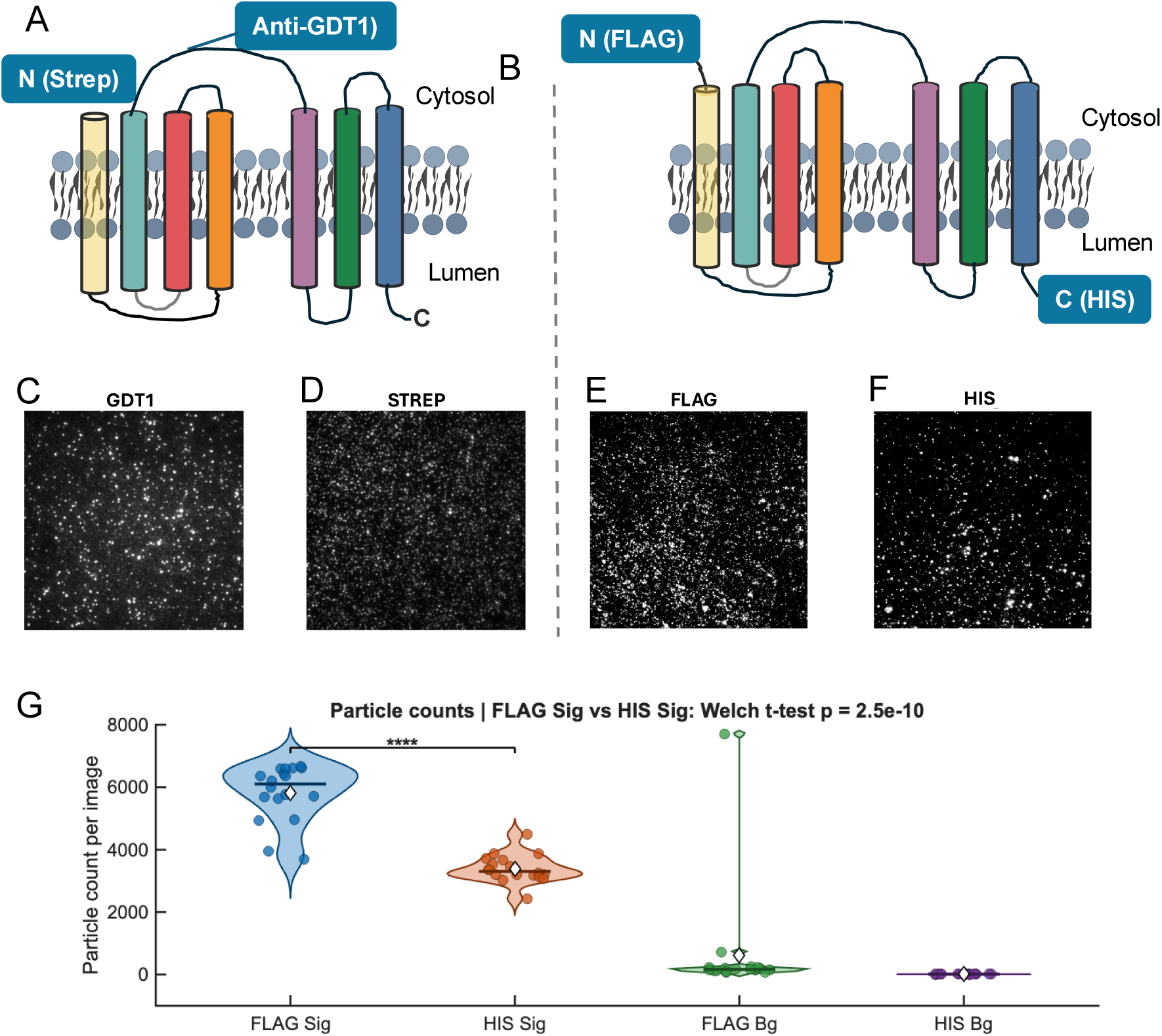
Antibody-accessibility measurements support membrane association and asymmetric epitope exposure of cell-free synthesized CaCA2 transporters in supported lipid bilayers. **(A,B)** Schematic representation of the expected SLB-associated geometries and antibody-accessible epitopes for **(A)** GDT1 and **(B)** TMEM165. For GDT1, anti-GDT1 antibody staining reports accessibility of the transporter, while C-terminal Strep-tag staining reports exposure of the C-terminal epitope. For TMEM165, N-terminal FLAG and C-terminal His tags were used to evaluate relative epitope accessibility in the SLB format. **(C, D)** Representative fluorescence microscopy images of GDT1-containing SLBs stained with **(C)** anti-GDT1 antibody or **(D)** anti-Strep antibody. **(E, F)** Representative fluorescence microscopy images of TMEM165-containing SLBs stained with antibodies against the **(E)** N-terminal FLAG tag or **(F)** C-terminal His tag. **(G)** Quantification of TMEM165 particle counts per image for FLAG signal, His signal, FLAG background, and His background controls. Background samples consisted of SLBs incubated with the corresponding fluorescent antibody in the absence of cell-free synthesized transporter. Each condition includes 18 images. Statistical comparison between FLAG and His signal was performed using Welch’s t test; **** indicates p < 0.0001.

We next evaluated the insertion and orientation of TMEM165 using antibodies against its N- and C-terminal epitopes. The predicted TMEM165 topology places the N-terminal FLAG tag and C-terminal His tag on opposite sides of the membrane (Fig. 4B). TIRF imaging showed detectable puncta for both epitopes, with a stronger signal observed for the FLAG epitope than for the His epitope (Fig. 4E, F). Quantification of 18 images collected across three independently prepared substrates showed approximately 6000 FLAG-positive puncta per image and approximately 3000 His-positive puncta per image, both above the corresponding background signals from SLBs lacking protein (Fig. 4G). The significantly higher FLAG signal compared with His signal indicates that most TMEM165 molecules adopt the orientation shown in Fig. 4B, while the lower but detectable His signal suggests the presence of a smaller population with opposite orientation, incomplete insertion, or partial C-terminal accessibility. Although differences in antibody affinity and epitope accessibility preclude quantitative comparison of the relative populations, the consistently greater FLAG signal supports a preferential orientation of TMEM165 corresponding to that shown in Fig. 4B.

Together with the low abundance of apparent truncation products observed by gel analysis, these results support cell-free synthesis and topology-consistent insertion of GDT1 and TMEM165 into SLBs. This antibody-accessibility assay therefore establishes the orientational context needed to evaluate transporter activity in the SLB-based platform.

To complement these measurements, activity was also assessed using the SLB platform, enabling direct, label-free measurement of ion transport. In this system, TMEM165 and GDT1 were synthesized directly into SLBs formed on PEDOT:PSS-coated electrodes (Fig. 5A). Ion transport was monitored using EIS, which detects changes in membrane electrical properties associated with ion flux across the bilayer. Following SLB formation and protein incorporation, impedance spectra were fit to an equivalent RC(RC) circuit, allowing extraction of membrane resistance (*R_m_*).^27, 28^ The intact bilayer produced a characteristic high-resistance “barrier” region in the impedance response (Fig. 5B). Upon addition of divalent cations, a decrease in this membrane resistance was observed, corresponding to ion transport through the embedded proteins. Specifically, the introduction of Ca^2+^ resulted in a clear drop in *R_m_*, indicating active transport across the membrane.

**Figure 5.**
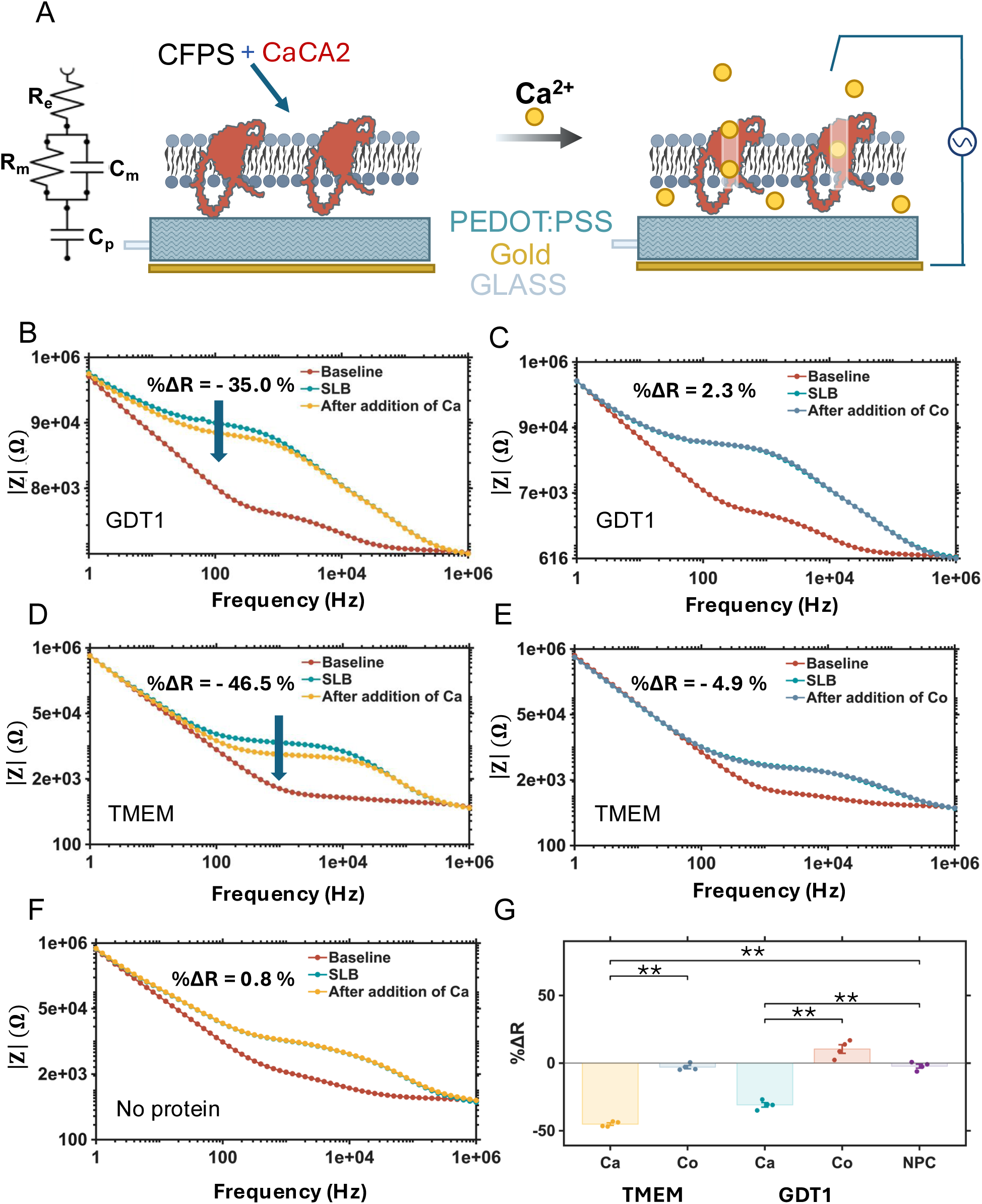
Functional characterization of GDT1 and TMEM165 in supported lipid bilayer platforms. (**A**) Schematic of the supported lipid bilayer (SLB)-based electrochemical platform and equivalent circuit model used to monitor membrane integrity and ion-dependent responses. The membrane is modeled as a parallel resistor–capacitor element, and membrane proteins are synthesized directly into the SLB prior to ion addition. (B,C) Representative Bode plots showing electrochemical impedance spectroscopy (EIS) responses of GDT1-containing SLBs following addition of (**B**) Ca²⁺ or (**C**) Co²⁺. (**D,E**) Representative Bode plots showing EIS responses of TMEM165-containing SLBs following addition of (**D**) Ca²⁺ or (**E**) Co²⁺. (**F**) Representative Bode plot of a protein-free SLB following ion addition, serving as a no-plasmid control. (**G**) Summary of EIS responses reported as percent change in membrane resistance (%ΔR) across all conditions. Statistical significance was determined using Welch’s t-test with Bonferroni correction (**p < 0.001, *p < 0.01, p < 0.05, NS, p > 0.05).

This approach further enabled direct comparison of ion-dependent responses across transporter and control conditions. GDT1-containing SLBs showed a 35.0% decrease in R_m_ following Ca²⁺ addition, whereas TMEM165-containing SLBs showed a 46.5% decrease (Fig. 5B,D). In contrast, Co²⁺ addition produced only minimal changes in R_m_ for both GDT1- and TMEM165-containing SLBs, with decreases of 2.3% and 4.9%, respectively (Fig. 5C, E). To determine whether the transport response extended to an additional divalent cation, we used Mn²⁺ shown in Fig S14. Mn²⁺ produced substantial decreases in membrane resistance for GDT1- and TMEM165 SLBs, with mean decreases of 33.5% and 39.7% respectively. This response was expected, as both GDT1 and TMEM165 have previously been reported to transport Mn²⁺. In contrast, protein-free SLBs showed a 10.4% increase in resistance following Mn²⁺ addition. Protein-free SLBs, prepared as a no-plasmid control (NPC), showed only a 0.8% change in R_m_ following ion addition with Ca²⁺ (Fig. 5F). Summary analysis of percent change in membrane resistance (%ΔR) demonstrated that Ca²⁺ produced significantly larger responses than Co²⁺ or NPC conditions (Fig. 5G), supporting that the observed resistance decreases arise from protein-dependent, ion-selective transport rather than nonspecific membrane disruption or electrode effects. Importantly, this platform allows interrogation of ion specificity without the need for encapsulated fluorescent reporters.

Together, these fluorescence- and EIS-based measurements demonstrate that GDT1 and TMEM165 are functionally active following cell-free synthesis into defined membrane environments and reflect native ion transport and known specificity. This validation is important because these proteins are complex, multi-pass intracellular transporters, and TMEM165 is normally produced within the cellular secretory pathway where folding, membrane insertion, and cellular processing may influence function. Despite the absence of most cellular processing pathways and post-translational modifications, TMEM165 and GDT1 retained measurable ion transport activity in this minimal system. The agreement between proteoliposome and SLB-based assays provides orthogonal validation of transporter activity and supports the use of this platform for mechanistic studies.

### Synthesis of TMEM disease associated variants

Moving beyond wild type proteins, the cell-free platform facilitates the study of clinically relevant targets as well. As such, the TMEM165 schematic highlights disease-associated mutations that have been identified in CDG patients (Fig. 6A, B). Domain coloring corresponds to the schematic in Fig. 6 A, B, facilitating comparison between topology and structure. The clinically observed variants are valuable not only because they are disease-linked, but because together they sample different modes of dysfunction such as direct perturbation of the conserved transport motif, indirect structural distortion of the transport pathway, and altered cellular localization or stability. Mapping these variants provides a framework for interpreting which defects are likely to arise from intrinsic changes in transport chemistry versus secondary effects that dominate in cells. For example, the E108G mutation (Patient 6) lies directly within the conserved acidic motif and is expected to perturb the electrostatic environment of the putative ion-coordination region.^29^ In contrast, R126H (Patient 4) and R126C (Patient 5) occur near the lysosomal targeting motif and are primarily linked to altered intracellular trafficking and Golgi depletion, that than direct catalytic disruption.^5^ G304R (Patient 5) lies in the seventh transmembrane region and has been proposed to indirectly distort the conformation of the acidic transport core despite being outside the canonical consensus motif.^30^ Here, this platform enables focused interrogation of TMEM165 variants and how disease-associated mutations govern intracellular ion transport.

**Figure 6.**
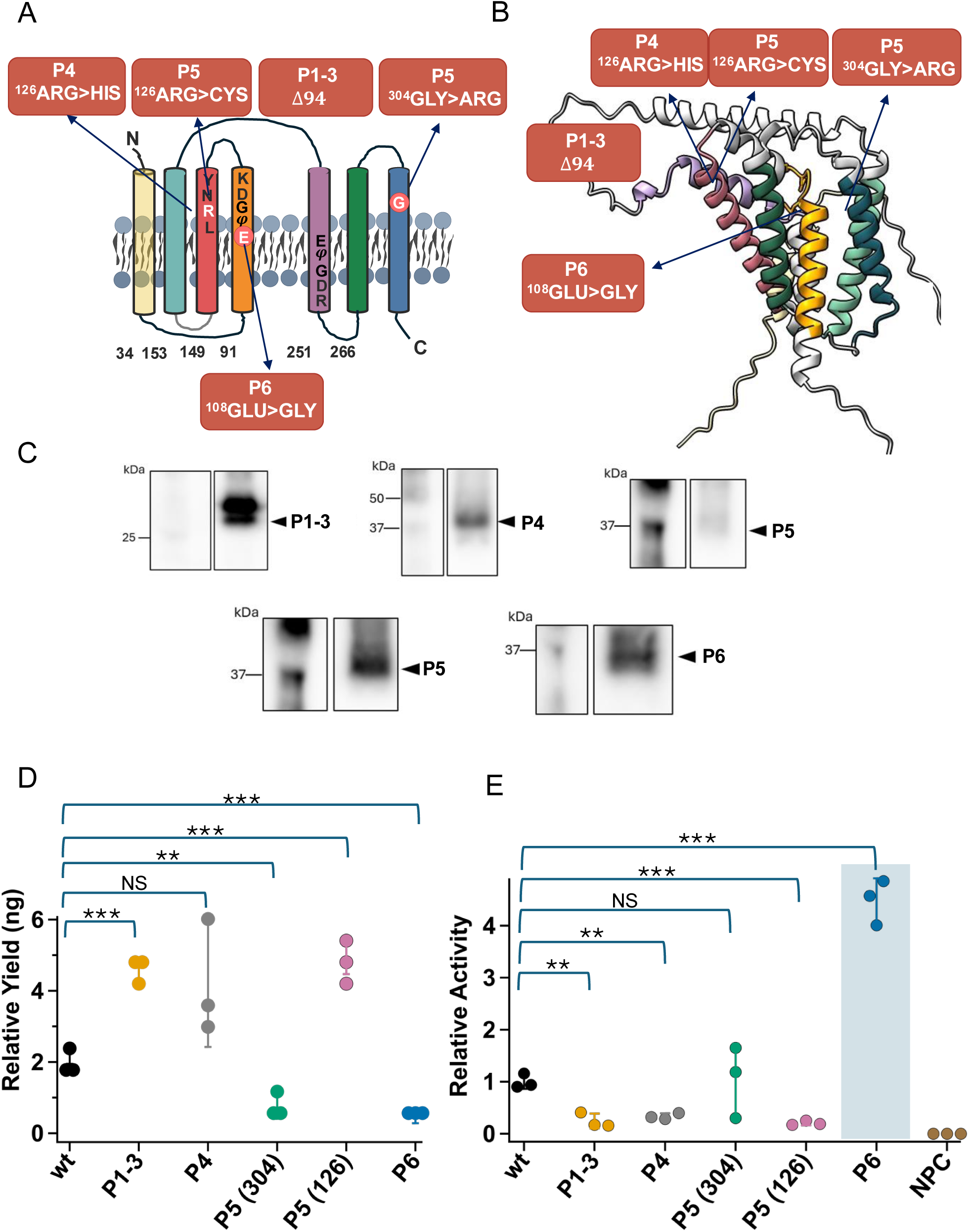
Clinically identified TMEM165 variants show altered expression and transport activity in the cell-free proteoliposome platform. **(A**) Schematic representation of TMEM165 membrane topology showing the locations of clinically identified mutations examined in this study, including E108G, R126H, R126C, G304R, and the 94-residue deletion associated with patients 1–3. **(B**) AlphaFold-predicted model of TMEM165 with the corresponding mutation sites mapped onto the predicted structure. **(C)** Western blot analysis of wild-type TMEM165 and clinically relevant TMEM165 variants following cell-free synthesis, with bands detected at the expected molecular weights**. (D**) Densitometric quantification of relative TMEM165 variant expression compared with wild-type TMEM165. **(E)** Relative transport activity of TMEM165 variants normalized to wild-type TMEM165. Statistical comparisons were performed between wild-type TMEM165 and each variant using Welch’s t test with Bonferroni correction. Significance is indicated as ***p < 0.001, **p < 0.01, *p < 0.05, and NS, p > 0.05. NPC, no-plasmid control.

To enable direct comparison of transport activity across TMEM165 variants, genes encoding wild type (wt) TMEM165 and five variants (Patients 1–3 (Δ94 internal truncation), Patient 4 (R126H), Patient 5 (R126C), and Patient 6 (E108G), along with G304R) were cloned into expression cassettes designed to minimize contributions from lac operator sequences and 5′ untranslated regions (UTRs), thereby improving translation efficiency in the cell-free system (Table S1). Cell-free expression was then used to produce wt and mutant TMEM165 proteins (Fig. 6C, D, S6-S8). To accurately quantify relative yields, a protein loading control (IF1, 10 kDa, Fig. S6) was included, allowing normalization of total protein levels across samples. Relative expression levels are shown in Fig. 6D.

Wt TMEM165 and the R126H mutant (Patient 4) exhibited comparable expression levels. In contrast, the Δ94 truncation (Patients 1–3) and the R126C mutant (Patient 5) showed increased apparent yields, while the G304R and E108G mutants exhibited reduced expression. These trends can be rationalized based on sequence and physicochemical considerations. However, while these mutations occur within the coding region of TMEM165 rather than regulatory elements such as the 5′ UTR, we cannot entirely exclude contributions from differences in transcription or mRNA stability in the cell-free system. The similarity in expression between wt and R126H is consistent with the comparable size and polarity of arginine and histidine. The increased yield observed for the Δ94 truncation likely reflects reduced energetic demand during translation due to the shorter polypeptide length. The R126C mutation may enhance expression by improving compatibility at the membrane interface; unlike arginine, which is bulky and positively charged, cysteine can better accommodate the membrane-solvent boundary and may reduce aggregation or ribosomal stalling during synthesis.^31^ In contrast, the reduced yields observed for G304R and E108G are likely influenced by codon usage and tRNA availability in the E. coli-based system. In particular, the E108G construct contains a glycine codon (GGG) that is translated less efficiently than other synonymous codons,^32^ while the introduction of arginine at position 304 may similarly introduce translational bottlenecks due to differences in tRNA abundance.^33^ These relative expression levels were used to normalize transport activity in subsequent analyses.

### Protein Activity of TMEM mutants

We next leveraged this reconstituted system to examine how clinically relevant TMEM165 mutations alter transport rate, ion selectivity, and pH-dependent regulation. All six mutant TMEM constructs were first assessed for total synthesis and membrane association relative to the wild-type TMEM165. Representative immunoblots presented in Fig. 6C, D, and Fig S7, S8 show varying steady-state levels of TMEM165 mutants produced during CFS reactions. Following successful expression of the TMEM165 mutants, transport activity was tested using the calcium reporter assay.

To compare activity across TMEM165 variants, fluorescence traces were first normalized to protein expression levels, as a direct relationship between relative yield and transport activity was observed (Fig. S9-S10). To further validate that fluorescence changes arise from transmembrane ion flux, diffusion was also benchmarked using the non-specific pore-forming protein α-hemolysin. At low protein concentrations, signal-to-noise was minimal; however, above a threshold concentration, fluorescence increased linearly with protein content, indicating that transport rate scales with the number of active pores (Figs S11–13). These results establish that the assay provides a quantitative readout of membrane permeability and can be used to relate protein concentration to activity.

The normalized fluorescence increase was then fit to a single-exponential model to extract apparent transport rates (Fig. S15). Comparison of relative activity across variants reveals distinct functional effects associated with each mutation (Fig. 6E). The Δ94 truncation (Patients 1–3), which removes a segment within the transmembrane region, results in reduced activity consistent with disruption of overall transporter architecture and loss of an intact transport pathway. The R126H mutation (Patient 4) also shows decreased activity, which may reflect altered electrostatic interactions near the membrane interface that influence ion access or local stability. The G304R mutation exhibits activity comparable to wild-type TMEM165, suggesting that although this substitution may perturb helix packing, it does not strongly disrupt transport under these conditions. In contrast, the R126C mutation (Patient 5) leads to decreased activity, potentially due to altered interfacial interactions or local structural destabilization. Notably, the E108G mutation (Patient 6), located within the conserved acidic motif, results in a substantial increase in apparent transport rate (∼4x). This enhancement may arise from loss of coordinated ion binding within the acidic region, effectively reducing steric contributions, allowing more rapid ion passage, and potentially reduced selectivity.

### Structural assessment of wild type versus E108G TMEM165

Structural models provide a framework for interpreting the altered transporter function of E108G and guiding the subsequent measurements. A closer view of the conserved acidic region (Fig. 7B) highlights residues Asp111, Glu108, Glu248, and Asp251, which have been proposed to contribute to divalent cation coordination. AMBER-based minimizations suggests that simultaneous coordination by all acidic residues is geometrically unfavorable, raising the possibility that ions interact with subsets of residues during transport. This interpretation is consistent with an alternating-access-type mechanism proposed for the UPF0016 family and with recent hypotheses of dynamic ion binding in structural studies.^5, 34, 35^ Figure 7A further highlights a flexible, acidic loop or “hinged” region proximal to the putative ion coordination site. This region contains additional charged residues and exhibits structural flexibility, suggesting a potential role in transient ion capture or electrostatic steering toward the transport pathway. While its precise function remains unresolved, its positioning and composition are consistent with a role in facilitating ion access to the core translocation region, as previously suggested.^34^ Substitution of the Glu108 mutant with glycine removes the negatively charged residue (Fig. 7C) within the putative transport pathway, suggesting that protonation state and transmembrane pH gradients may play a role in modulating TMEM165 activity. ^36, 37^ Together, these structural features motivate a model in which electrostatic architecture and residue-specific perturbations govern ion coordination and transport behavior.

**Figure 7.**
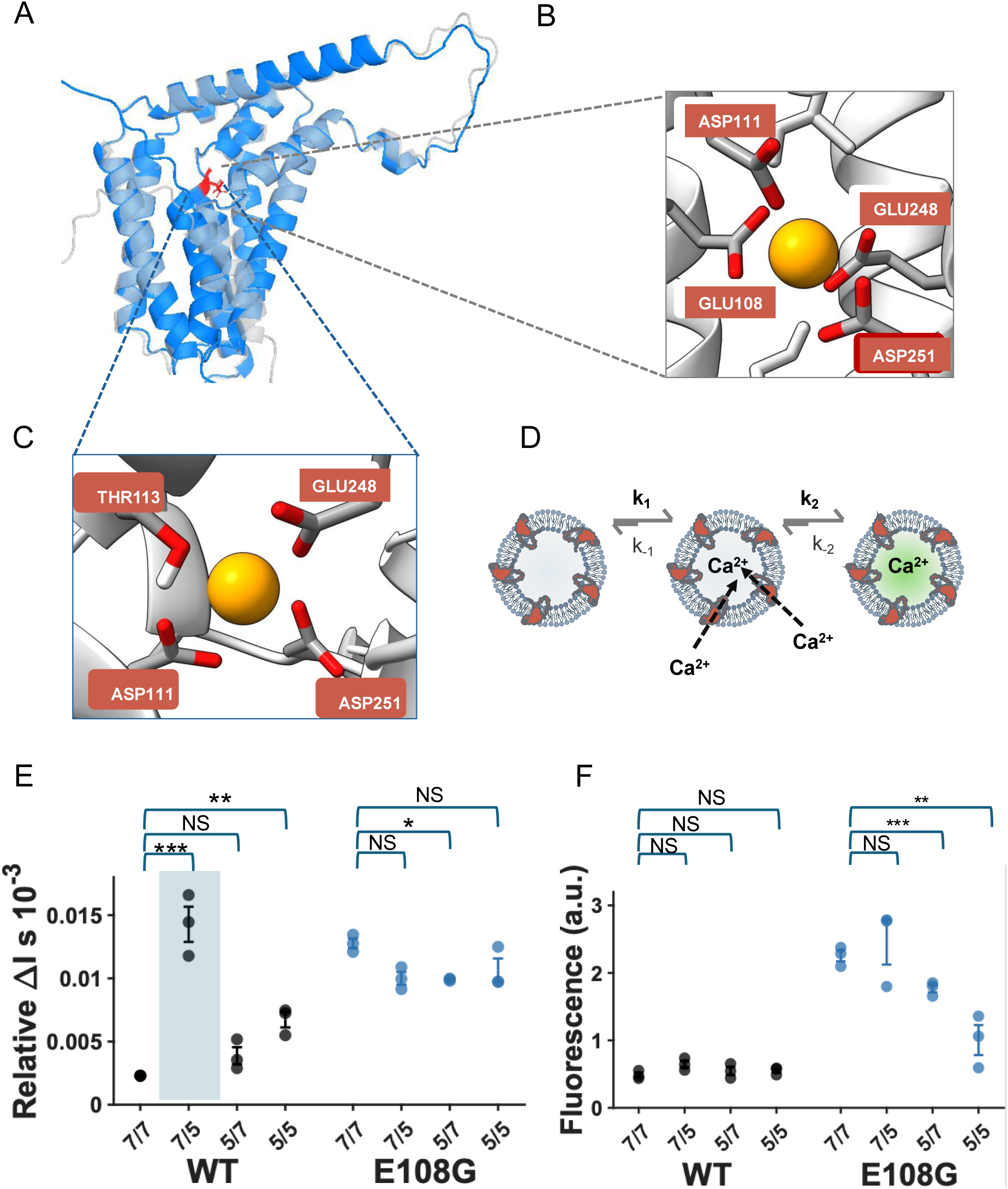
Structural modeling and pH-dependent transport measurements reveal altered Ca²⁺ transport behavior in the TMEM165 E108G variant. **(A)** Overlay of AlphaFold-predicted structures of wild-type TMEM165 and the E108G variant, shown in light blue and dark blue, respectively. The putative acidic ion-coordination region is highlighted in red. **(B)** Structural model of TMEM165 with a docked Ca²⁺ ion, highlighting the putative ion-coordination region formed by acidic residues surrounding the predicted binding site. **(C)** Magnified view of the putative ion-coordination region in the E108G variant, showing the local environment surrounding residue 108 after substitution of glutamate with glycine. (**D**) Kinetic scheme for the fluorescence-based Ca²⁺ transport assay. In this model, k₁ and k₋₁ represent Ca²⁺ movement across the proteoliposome membrane, whereas k₂ and k₋₂ represent Ca²⁺ binding and unbinding to the encapsulated fluorescent sensor. Under these assay conditions, membrane transport was treated as the rate-limiting step. (**E**) Relative transport rates of wild-type TMEM165 and E108G measured under defined pH gradients (pH_in_/pH_out_): 7/7, 7/5, 5/7, and 5/5, with values normalized to relative protein expression. (**F**) Steady-state fluorescence responses measured under the same pH conditions and normalized to Triton X-100 permeabilization to account for the maximum sensor-accessible signal. Statistical comparisons were performed using Welch’s t test with Bonferroni correction. Significance is indicated as ***p < 0.001, **p < 0.01, *p < 0.05, and NS, p > 0.05.

### Interrogating the Effect of H^+^ Transmembrane Gradients on Transporter Activity

To probe the mechanistic basis of the altered transport behavior observed for the E108G mutant (Fig 6E), we examined the role of pH, given that this mutation lies within the conserved acidic motif implicated in ion coordination and electrostatic regulation. To test this, calcium transport was evaluated under defined transmembrane pH conditions using fluorescence-based measurements, which provide access to transport kinetics on experimentally relevant timescales (Fig. 7D, S16, S17). To probe both proton dependence and ion selectivity, we compared transport rates for wild-type and E108G under defined transmembrane pH conditions, reported as pH***x****_in_*/pH***y****_out_*, where pH***x****_in_* corresponds to the intraliposomal environment and pH***y****_out_* to the external buffer. Four conditions were tested: 7/7, 5/5, 7/5, and 5/7. The selected pH conditions (7 and 5) were chosen to approximate physiologically relevant environments encountered by TMEM165, spanning near-neutral cytosolic conditions (pH ∼7) and the acidic luminal conditions of intracellular organelles, such as the Golgi and endosomal compartments (pH ∼5–6). Of these conditions, 5/7 most closely reproduces the physiological orientation encountered by TMEM165 across acidic organellar membranes, with the intraliposomal compartment representing the acidic lumen and the external buffer representing the near-neutral cytosol; pH 5 is particularly representative of the lysosomal lumen, whereas the Golgi lumen is generally somewhat less acidic.

Across all conditions, the E108G mutant exhibited relatively constant transport (still elevated with respect to wild type), showing little sensitivity to pH (Fig. 7E). In contrast, wild-type TMEM165 displayed strong pH dependence. The highest activity (∼7x fold increase) was observed under the 7/5 condition, followed by 5/5 (∼3 - 4x fold increase) and 5/7 (∼2x fold increase), all relative to the 7/7 condition. These results indicate that wild-type activity is strongly modulated by transmembrane proton gradients, whereas this responsiveness is largely lost in the E108G mutant.

In CaCA2 transporters, divalent cation transport is thought to be coupled to proton exchange, such that protonation of acidic residues within the transport pathway modulates ion binding affinity and translocation kinetics. Thus, transmembrane pH gradients can directly influence transport by altering the protonation state of coordinating residues and the electrostatic landscape of the pathway.

To interpret these differences in activity, we examined structural models of the conserved acidic region (Fig. 7B, C). In wild-type TMEM165, residues within this “acidic cage” are positioned to coordinate divalent cations and participate in proton-dependent transport. Substitution of Glu108 with glycine removes a key negatively charged residue located near the solvent-accessible entrance of the ion transport pathway. While modeling suggests that Ca²⁺ coordination may be partially retained through neighboring residues, the loss of this acidic group disrupts the local electrostatic environment and eliminates a key pH-sensitive interaction. As a result, the mutant retains transport activity but loses sensitivity to proton gradients, consistent with the observed steady activity across conditions.

The behavior of wild-type TMEM165 further supports a role for proton activated transport.^36^ The elevated activity under the 7/5 condition (acidic external environment) is consistent with proton activation or proton-assisted transport. While a high apparent rate under asymmetric external acidification may initially seem counterintuitive for a putative exchanger, this observation strongly aligns with recent findings,^36^ which demonstrated that TMEM165 functions as a proton-activated importer where cytosolic acidification serves as a requisite switch to trigger transport. Although the 7/5 condition is unlikely to reflect the physiological pH gradient experienced by TMEM165 in vivo, the strong response under this condition further supports a role for proton activation, even if such activation occurs over a less extreme pH range physiologically. The 5/5 condition yields the second highest activity, suggesting that global acidification enhances transport, potentially through increased protonation of key residues (E248 and D251) coupled with proton activation on the outside. The 5/7 condition, which more closely reflects gradients between acidic organelles and the cytosol, results in moderate activity, indicating that internal acidification alone is insufficient to maximize transport. Notably, under the 7/5 condition, wild-type activity approaches that of E108G, suggesting that protonation of residues in the wild-type protein may partially mimic the reduced electrostatic barrier introduced by the mutation.

This observation can also be rationalized by comparison of the TMEM165 pore region and relative electrostatic surface potential, which suggest that the E108G mutant has a wider pore entrance and reduced negative surface potential due to loss of a critical negatively charged residue near the pore entrance (Fig. 8 A, B, C). It is important to note that increased transport rate does not necessarily correspond to improved function in cells. Intracellular transporters such as TMEM165 operate to maintain ion homeostasis, and perturbations that increase flux or reduce selectivity may disrupt the balance of ion gradients required for proper organelle function.

**Figure 8.**
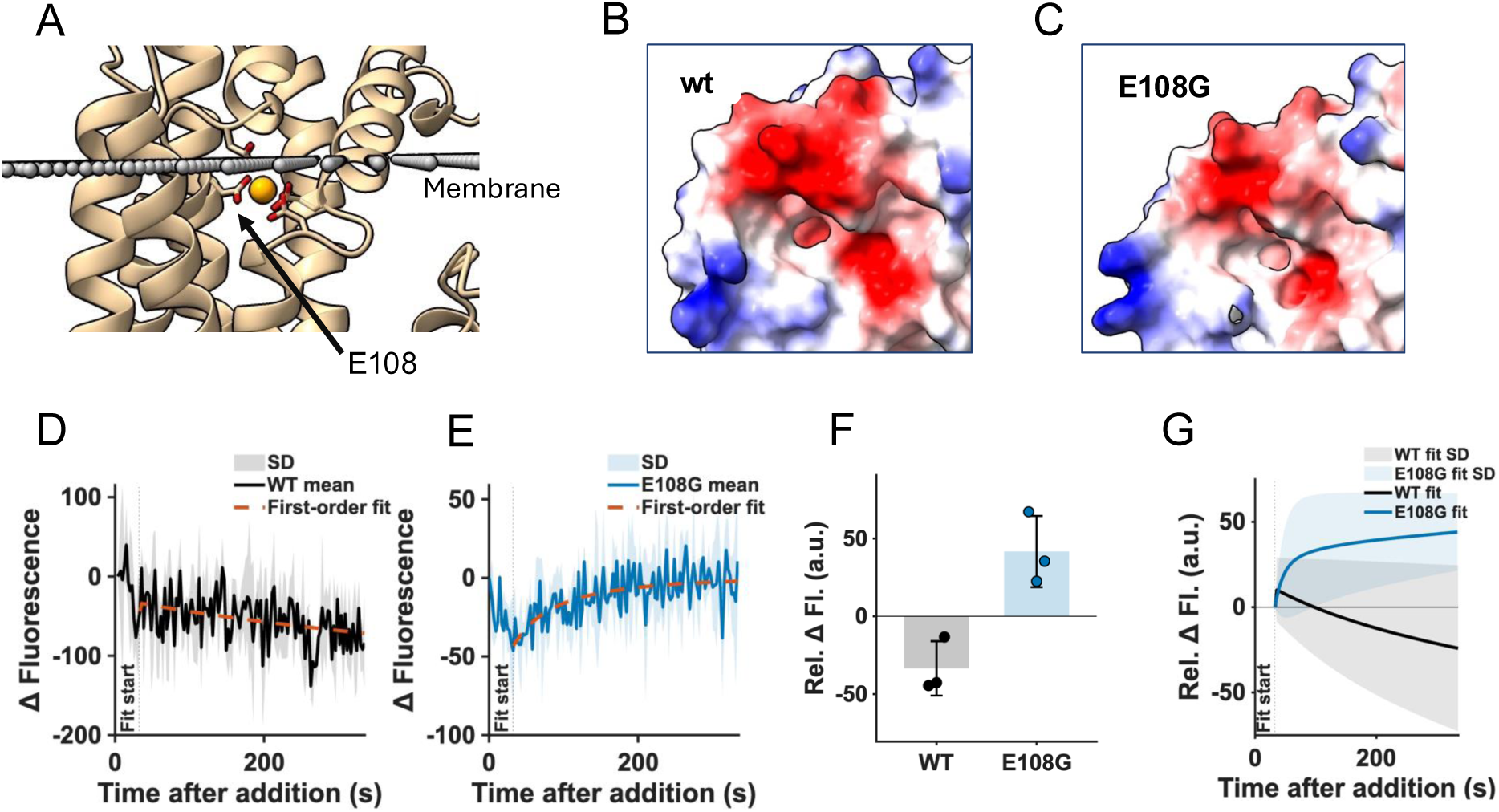
The E108G mutation alters the local electrostatic environment and changes sodium permeability in TMEM165 proteoliposomes. (**A**) Structural model of TMEM165 with a docked Ca²⁺ ion following Amber minimization and membrane positioning in a DOPC bilayer using the PPM server. The membrane plane and residue E108 are indicated. (**B, C**) Electrostatic surface potential maps of the putative ion-binding region in (**B**) wild-type TMEM165 and (**C**) E108G TMEM165. Red indicates negative surface potential and blue indicates positive surface potential. (**D, E**) Representative fluorescence traces from proteoliposomes containing encapsulated sodium sensor and either (**D**) wild-type TMEM165 or (**E**) E108G TMEM165 following external sodium addition. Changes in fluorescence report sodium movement into the liposomal lumen. Dashed lines indicate first-order fits to the initial response after sodium addition. (**F**) Quantification of steady-state fluorescence change for wild-type TMEM165 and E108G across three independent experiments. (**G**) Direct comparison of first-order fits from wild-type TMEM165 and E108G sodium transport traces. Shaded regions indicate the standard deviation across three independent experiments. Statistical comparisons were performed using Welch’s t test with Bonferroni correction.

We also compared steady-state behavior under these conditions. For wild-type TMEM165, despite substantial differences in initial rates, the steady-state fluorescence signal was similar across all pH conditions (Fig. 7F), indicating convergence to a comparable equilibrium. In contrast, the E108G mutant exhibited condition-dependent steady-state behavior. While the 7/7 and 7/5 conditions were similar to wild-type, the steady-state signal decreased under the 5/7 condition and decreased further under the 5/5 condition. This difference between initial rate and steady-state behavior suggests that the mutation alters both transport kinetics and equilibrium properties. One possible explanation is that wild-type TMEM165 maintains a coupled transport mechanism, potentially involving coordinated exchange of Ca²⁺ and protons, allowing the system to reach a consistent steady state despite differences in the initial rates. In contrast, disruption of the acidic residue in E108G may impair this coupling, leading to altered occupancy of the transport pathway or reduced ability to sustain ion exchange under certain pH conditions. As a result, although the mutant exhibits higher initial rates, it ultimately transports fewer ions at equilibrium under specific conditions, consistent with a loss of coupling and selectivity. Consistent with this interpretation, we examined whether TMEM165 transport is accompanied by changes in intra-liposomal pH. Liposomes were loaded with the pH-sensitive dye FITC, and its responsiveness was first validated over the relevant pH range (pH 5–7) (Fig. S18). When liposomes were prepared with an internal pH of 5 and exposed to an external pH of 7, wild-type TMEM165 exhibited a gradual increase in internal pH over time, consistent with proton efflux coupled to ion transport (Fig. SI 18). In contrast, the E108G mutant showed little to no change in internal pH, similar to control liposomes lacking protein. These results provide evidence that proton coupling is preserved in the wild-type protein but is disrupted in the mutant, supporting the interpretation that E108G alters the coupling mechanism underlying transport.

To further examine ion selectivity, we performed complementary experiments using a sodium-sensitive fluorescent reporter. Under identical conditions, wild-type TMEM165 showed no measurable Na⁺ transport (Fig. 8E, G, H, S19), whereas the E108G mutant produced a clear increase in signal, indicating sodium permeability (Fig. 8F, G, H). This loss of selectivity is consistent with disruption of the conserved acidic residue, which likely contributes to defining the electrostatic constraints that restrict ion passage. Together with the pH measurements, these results suggest that the E108G mutation weakens coupling ion transport from proton gradients and relaxes substrate specificity, allowing passage of ions that are normally excluded. Such altered ion permeability may have implications for charge balance across the membrane, a possibility that will require direct membrane-potential measurements in future studies.

## Discussion

In this work, we establish a minimal, cell-free membrane platform for dissecting the intrinsic transport mechanisms of intracellular ion transporters under defined biochemical and electrochemical conditions. By combining cell-free synthesis with reconstituted liposome and SLB systems, we directly controlled ionic conditions, pH gradients, and protein identity while avoiding the confounding effects of trafficking, organelle composition, compensatory ion regulation, and cellular homeostasis. Within this framework, we show that ion transport in TMEM165 is governed by an electrostatic network that couples divalent cation movement to transmembrane proton gradients. Importantly, this platform enabled direct comparison of wild-type TMEM165 with a disease-associated variant, revealing how a clinical mutation alters the underlying transport mechanism.

The inclusion of GDT1 provided a mechanistically defined reference point for interpreting TMEM165 function. As a member of the same CaCA2 family with prior characterization in model organisms, GDT1 offered a tractable system for establishing expected transport behavior in a minimal membrane context. The observation of consistent transport activity and ion selectivity for both GDT1 and TMEM165 indicates that the platform preserves intrinsic functional features of this transporter family, thereby supporting direct mechanistic interrogation of TMEM165 and its variants.

To further dissect transporter behavior, we leveraged the complementarity of the two membrane systems. Liposome-based assays provided access to transport kinetics through fluorescence readouts, enabling quantification of rates under defined ionic and pH conditions. In contrast, the SLB platform coupled with EIS provided a direct, label-free measurement of ion-dependent membrane responses and allowed interrogation of ion selectivity without encapsulated reporters. These approaches probe distinct but complementary transporter functions: optical assays resolved transport dynamics, kinetic responses, and luminal ion accumulation, whereas EIS provided an orthogonal electrical readout of membrane-associated ion transport. Together, these methods strengthen interpretation of transport mechanisms by reducing reliance on a single readout.

Using this framework, we observed that the E108G mutation produced a substantial increase in apparent transport rate relative to wild-type TMEM165. Because this mutation removes a conserved acidic residue within the proposed ion coordination region, the increase in activity is unlikely to reflect enhanced function. Instead, it suggests disruption of electrostatic interactions that normally regulate transport. By directly comparing wild-type and disease-associated TMEM165 variants in a controlled membrane environment, we found that disruption of E108 increased apparent transport rates, decreased ion specificity, and markedly reduced pH sensitivity. Although this finding contrasts with the reduced activity reported for E108G in cellular systems,^30^ the difference may reflect faster but less selective or less effectively coupled ion flux in the reconstituted membrane environment. These results indicate that transport rate alone is not a sufficient measure of function and instead highlight the importance of electrostatic coupling in maintaining ion selectivity and gradient responsiveness.

Taken together, the results support a model in which the electrostatic environment of the conserved acidic region governs both ion selectivity and coupling to proton gradients. In the wild-type protein, coordinated interactions between acidic residues and permeating ions likely enforce selectivity and enable proton-coupled ion movement. Our results support that TMEM165 functions as an antiporter, in which Ca²⁺ influx is coupled to proton efflux. Additionally, TMEM165 is proton activated, with transport rates increasing upon E108 protonation. In a biological context, E108 protonation likely occurs as protons are effluxed, promoting conformational changes that allow Ca²⁺ influx. Because our system allows us to sample environments that are not readily accessible in cells, we were able to alter the protonation state of this residue using the exterior environment, rather than only through the antiporter mechanism. Mutation of Glu108 appears to disrupt this electrostatic network, lowering the barrier for ion passage while weakening coupling to proton gradients. This leads to increased apparent transport rates but decreased selectivity and altered equilibrium behavior.

These findings are derived from a minimal, cell-free system that isolates intrinsic transporter behavior from cellular complexity. While additional factors in vivo, including membrane composition, accessory proteins, ion buffering systems, and trafficking pathways, will further modulate function, the controlled framework used here enables direct resolution of the physicochemical principles governing ion transport. In this context, the differences observed between wild-type and mutant proteins reflect fundamental changes in transport mechanism that are likely to underlie their cellular phenotypes. The platform is therefore particularly useful for studying clinically relevant variants whose cellular effects may combine changes in expression, localization, stability, and intrinsic transport activity.

This work establishes a generalizable framework for studying intracellular membrane proteins using minimal, reconstituted systems. By integrating complementary optical and electrical measurements, we provide evidence that electrostatic architecture governs ion transport, selectivity, and proton coupling in TMEM165, and show how disease-associated mutations disrupt these relationships. More broadly, this approach enables systematic deconvolution of membrane protein function under controlled conditions, providing a route to resolve mechanisms that are otherwise obscured in complex cellular environments. For TMEM165, this framework reveals that the clinically relevant E108G mutation perturbs the electrostatic coupling network required for selective, pH-responsive antiport, highlighting how minimal membrane platforms can connect molecular mechanism to disease-associated transporter dysfunction.

## Materials and Methods

### Plasmids and DNA constructs

All plasmids used in this study contained a T7 promoter and ribosome-binding sequence to enable expression with the T7 polymerase-based PUREfrex 2.1 cell-free protein synthesis system. Different construct sets were used for synthesis tracking, orientation measurements, and functional transport assays. For fluorescence-based tracking of protein synthesis and membrane association, full-length GDT1-sfGFP and full-length TMEM165-sfGFP constructs were used. These constructs were generated in pET28a-based vectors by Twist Bioscience and contained a C-terminal superfolder GFP tag. A TEV protease cleavage site was placed between the transporter and sfGFP so that GFP removal could be used to evaluate protease accessibility after membrane insertion. For functional measurements of the original full-length transporters, non-GFP-tagged GDT1 and TMEM165 constructs were used. These constructs were used for initial calcium transport assays, supported lipid bilayer measurements, and comparison of GDT1 and TMEM165 activity. For the TMEM165 patient-variant transport experiments, the matched wild-type TMEM165 construct and all TMEM165 disease-associated variants were generated in the pMA-RQ vector backbone by GeneArt. This set included wild-type TMEM165, the Δ94-residue truncation associated with patients 1-3, R126H, R126C, G304R, and E108G. These constructs were used as a matched set for expression comparison, western blot quantification, and proteoliposome transport assays. The use of the same pMA-RQ backbone for the wild-type and variant TMEM165 constructs allowed direct comparison across the variant series.

### Liposome preparatio

Liposomes were prepared from chloroform stocks of phospholipids. For standard calcium transport assays, DOPC liposomes were prepared from 1,2-dioleoyl-sn-glycero-3-phosphocholine 18:1 PC. Lipids were dried under a stream of nitrogen for at least 5 min to form a thin lipid film and then placed under vacuum for at least 3 h to remove residual chloroform. Dried lipid films were hydrated with the appropriate internal buffer containing the fluorescent reporter (150 mM Tris-HCl, 150 mM KCl, pH 7.4 for EIS or 50 mM Tris, 150 mM NaCl, 0.1 mM EGTA, pH 7.4 for fluorescent assays). Fluorescent reporter (Fluo-4 FF) was prepared in the fluorescent assay buffer at 50 µM. Unless otherwise indicated, liposomes used for fluorescence-based transport assays were prepared at a final lipid concentration of 26.09 mM. For experiments that required stable transmembrane pH gradients, including pH-dependent calcium transport assays and FITC-based intraliposomal pH measurements, a different lipid formulation was used. These liposomes were prepared from a DOPC/POPC mixture containing 15 mol% cholesterol. This formulation was used because DOPC-only liposomes ruptured or became unstable during exposure to different internal and external pH conditions. The cholesterol-containing formulation improved liposome stability during pH exchange and allowed the pH-gradient experiments to be performed reproducibly. After hydration, liposome suspensions were subjected to one freeze-thaw cycle consisting of 5 min at -80°C followed by 5 min at room temperature. Samples were vortexed after thawing and extruded at least 29 times using an Avanti Mini-Extruder. Liposomes used for solution-based transport assays were extruded through 200 nm Whatman Nuclepore polycarbonate membranes. Liposomes used for supported lipid bilayer formation were extruded through 100 nm membranes. Reporter-loaded liposomes were purified by gravity size-exclusion chromatography using Sepharose columns to remove unencapsulated dye. For each preparation, 100 µL of liposome suspension was applied to the column, and the liposome-containing fraction was collected for downstream cell-free protein synthesis or transport measurements.

### Liposome characterization

Liposome size was measured by dynamic light scattering using a Zetasizer Nano-ZS instrument equipped with a 4 mW He-Ne laser operating at 632 nm and a backscatter detector angle of 173°. Measurements were collected after extrusion and, when relevant, after size-exclusion chromatography. DOPC liposomes used for standard calcium transport assays were approximately 178 nm in diameter after extrusion through 200 nm membranes and approximately 172 nm after SEC purification. Liposomes used for supported lipid bilayer formation were prepared by extrusion through 100 nm membranes before bilayer assembly.

### Cell-free protein synthesis into liposomes

Cell-free protein synthesis was performed using the PUREfrex 2.1 Cell-Free Protein Synthesis Kit according to the manufacturer’s protocol. Reactions were assembled with 400 ng plasmid DNA per 20 µL reaction volume. For synthesis into liposomes, plasmid DNA, PUREfrex reaction components, and preformed liposomes were mixed directly. Unless otherwise indicated, liposomes were present at a final lipid concentration of 8.7 mM. Reactions were incubated at 37°C for 4 h. Protein synthesis was mostly complete within 2-3 h but 4 hours was suggested by manufacturer. Proteoliposomes and controls were kept at 4°C before experiments. For experiments requiring purified proteoliposomes, samples were passed through size-exclusion columns before analysis.

### Cell-free protein synthesis into supported lipid bilayers

Supported lipid bilayers were formed before addition of the cell-free reaction components. For fluorescence microscopy and antibody-accessibility measurements, DOPC liposomes were used to form bilayers on glass supports. For electrochemical impedance spectroscopy measurements, POPC liposomes were used to form bilayers on PEDOT:PSS-coated gold electrodes. PDMS wells were prepared from Sylgard 184 using a 10:1 elastomer-to-crosslinker ratio and attached to the substrate surface. Each well had an approximate area of 1 cm². Liposome solutions were added to the wells and incubated for 20-45 min to allow vesicle adsorption and rupture. For glass-supported bilayers, 1 mg/mL DOPC liposomes were used. For electrode-supported bilayers, 2 mg/mL POPC liposomes were used. After bilayer formation, wells were rinsed with buffer (125 mM HEPES, 150 mM KCl pH 7.4) to remove excess unruptured liposomes. For direct synthesis into SLBs, PUREfrex reaction components were assembled without additional liposomes and applied directly onto the preformed bilayer. Reactions were incubated on the SLB surface at 37°C for 30 min unless otherwise stated. After incubation, wells were rinsed with buffer (150 mM Tris-HCl, 150 mM KCl, pH 7.4) to remove soluble reaction components and stop the reaction.

### PEDOT:PSS/gold electrode preparation

Gold microelectrodes were fabricated on fused silica substrates using photolithography. Gold contact pads were first patterned, followed by deposition of an insulating SiO₂ layer. A second lithography step was used to define the electrode openings and expose the underlying gold. PEDOT:PSS containing 1% GOPS was spin-coated onto the patterned substrates and baked at 140°C for 30 min. A final patterning step was used to remove PEDOT:PSS from the insulating regions, leaving PEDOT:PSS only on the electrode areas. Before use, completed PEDOT:PSS/gold electrode devices were soaked in 100 µM KCl for more than 24 h. Immediately before SLB formation, electrode surfaces were treated with oxygen plasma for 30 s using a Harrick Plasma cleaner. POPC liposomes were then added to the plasma-treated surface for SLB formation.

### Western blot analysis

Cell-free expressed proteins were analyzed by SDS-PAGE and western blotting. Samples were separated on 10-12% polyacrylamide gels for 45 min at 175 V. Proteins were transferred to PVDF membranes by wet transfer for 1 h at 20 V. Membranes were blocked for 1 h at room temperature in 5% milk in TBST buffer containing 50 mM Tris, 150 mM NaCl, and 0.5% Tween-20, pH 7.6. Membranes were then incubated overnight at 4°C with primary antibody diluted in TBST. His-tagged proteins were detected using mouse anti-6xHis antibody diluted 1:2000. GFP-tagged constructs and TEV cleavage products were detected using anti-GFP antibody. After primary antibody incubation, membranes were washed three times for 5 min each in TBST. Membranes were then incubated for 1 h at room temperature with HRP-conjugated secondary antibody diluted 1:5000 in TBST. After secondary antibody incubation, membranes were washed three times for 5 min each and developed using Clarity Western ECL Substrate before imaging. Antibody dilutions were all per manufacturer’s suggestions. Band intensities were quantified by densitometry. For TMEM165 variant expression comparisons, relative protein levels were normalized to the IF1 loading control. These normalized expression values were used to correct transport activity across the variant series.

### TEV protease cleavage assay

TEV protease cleavage was used to assess accessibility of the C-terminal sfGFP tag after membrane insertion. For proteoliposome samples, TEV protease was added to SEC-purified proteoliposomes containing full-length transporter-sfGFP constructs. Samples were incubated at 30°C in the dark to limit GFP photobleaching. Fresh TEV protease (New England Biolabs) was added every hour for 3 h where indicated with dilutions as per manufacturers protocol. Samples were then analyzed by western blotting using anti-GFP antibody. The appearance of a cleaved GFP-containing band was used as evidence that the TEV site was accessible to protease. For SLB samples, TEV protease was added directly to supported lipid bilayers containing cell-free synthesized GFP-tagged transporters. Samples were incubated at 30°C in the dark for 1 hour, and total internal reflection fluorescence microscopy was used to image GFP fluorescence before and after TEV treatment. Loss of membrane-associated GFP fluorescence after TEV addition was used as a readout of protease accessibility in the planar bilayer format.

### Antibody-accessibility assay in supported lipid bilayers

Protein insertion and orientation in SLBs were assessed using antibody-accessibility measurements. After cell-free synthesis into SLBs, samples were blocked with 0.1 mg/mL BSA for 20 min at room temperature. SLBs were then incubated with primary antibody for 30 min, rinsed with buffer, and incubated with fluorescent secondary antibody for 30 min. After a final rinse, samples were imaged by TIRF-M. For GDT1, an anti-GDT1 antibody was used to detect the predicted exposed loop region, and an anti-Strep antibody was used to assess accessibility of the C-terminal Strep tag. For TMEM165, anti-FLAG and anti-His antibodies were used to detect the N-terminal FLAG and C-terminal His epitopes, respectively. All antibody concentrations were in accordance with manufacturers protocols. Background controls consisted of SLBs treated with the same antibody conditions in the absence of cell-free synthesized transporter.

### TIRF microscopy and particle-count analysis

SLB-associated fluorescence was imaged using TIRF-M. Images were collected using identical acquisition settings within each experiment so that particle counts could be compared across conditions. For particle-count analysis, images were processed in MATLAB. Raw images were background corrected, smoothed or contrast enhanced as indicated, and thresholded using the same analysis parameters within each experimental set. Fluorescent puncta were identified as connected components within a defined size range, and particle counts were reported as the number of detected puncta per image. For TMEM165 antibody-accessibility measurements, 18 images were analyzed per condition across three independently prepared substrates. Conditions included FLAG signal, His signal, FLAG background, and His background. Statistical comparisons were performed using Welch’s t test.

### Calcium transport assay in proteoliposomes

Calcium transport was measured using proteoliposomes containing encapsulated Fluo-4FF. DOPC liposomes were loaded with Fluo-4FF, purified by size-exclusion chromatography (SEC), and used for cell-free synthesis of GDT1, TMEM165, or TMEM165 variants. Final concentration of encapsulated reporter was calculated from the SEC profile in Fig. S1 and was 3-5% of total Fluo-4 FF added to the initial buffer, therefor the final concentration of Fluo-4 FF added to the cell-free solutions was 1.5-2.5 µM and 0.04-0.075 µM in the calcium assay. For each measurement, 27 µL of assay buffer containing 50 mM Tris, 150 mM NaCl, and 0.1 Mm EGTA, pH 7.4, was mixed with 3 µL of cell-free proteoliposome sample in a black 384-well plate. Baseline fluorescence was recorded for 5 min using a Tecan Spark plate reader with excitation and emission wavelengths of 464 nm and 518 nm, respectively. After baseline acquisition, 5 µL of calcium-containing buffer containing 50 mM Tris, 150 mM NaCl, 0.1 mM EGTA, and 4 mM CaCl₂, pH 7.4, was injected into the well. Calcium transport was monitored immediately by recording fluorescence at 1 s intervals for at least 15 min. At the end of each experiment, 3 µL of 30% Triton X-100 was added to solubilize the liposomes and release all encapsulated reporter. Fluorescence changes were normalized to the maximum fluorescence measured after Triton X-100 addition. The linear response range of Fluo-4FF was established before transport measurements by measuring fluorescence as a function of calcium concentration. Assay conditions were selected so that external calcium was in excess and the encapsulated sensor remained below saturation. Protein-free liposomes were used as negative controls to confirm that the observed fluorescence increase required transporter expression.

### pH-gradient calcium transport assays

To determine how transmembrane pH gradients affect TMEM165 transport, calcium transport was measured under defined internal and external pH conditions. These experiments were performed using DOPC/POPC liposomes containing 15 mol% cholesterol because this formulation maintained liposome integrity during pH exchange. Liposomes were prepared with internal buffer at pH 5 or pH 7 (citrate or tris buffer respectively), purified by SEC, and then diluted into external buffer at pH 5 or pH 7 immediately before measurement. Four pH conditions were tested: pH_in_/pH_out_ 7/7, 7/5, 5/7, and 5/5. Proteoliposomes containing wild-type TMEM165 or E108G TMEM165 were measured using the same Fluo-4FF calcium transport assay described above. Calcium transport was initiated by external calcium addition, and fluorescence was recorded immediately. Initial fluorescence increases were fit to a single-exponential model, and apparent transport rates were normalized to relative protein expression. Steady-state fluorescence values were normalized to the Triton X-100 maximum signal to account for differences in total encapsulated reporter.

### Sodium transport assay

Sodium permeability was assessed using proteoliposomes containing encapsulated ING-2 TMA⁺ salt, a membrane-impermeable sodium-sensitive fluorescent reporter. ING-2 was used at a working concentration of 3 µM in 20 mM HEPES, 100 mM KCl, pH 7.45. ING-2-loaded liposomes were prepared, purified, and used for cell-free synthesis as described for the calcium transport assay. Wild-type TMEM165 and E108G TMEM165 were compared under identical conditions. After baseline acquisition, sodium transport was initiated by addition of 4 µL of 0.2 M NaCl stock to the external buffer, and fluorescence was monitored immediately using a plate reader. ING-2 fluorescence was measured at an emission wavelength of 545 nm. Initial response phases were fit to a single-exponential function, and steady-state fluorescence changes were quantified across independent experiments. Protein-free liposomes were used as controls. Sodium transport was interpreted from the increase in ING-2 fluorescence following external sodium addition.

### FITC-based intraliposomal pH assay

Changes in intraliposomal pH were monitored using FITC-loaded liposomes. These experiments were performed using DOPC/POPC liposomes containing 15 mol% cholesterol to preserve liposome stability during exposure to different internal and external pH conditions. FITC was prepared as a 100 mg/mL stock solution in DMSO and diluted 1:1000 into the liposome hydration buffer. Liposomes were hydrated in the FITC-containing buffer at the desired internal pH and then purified by size-exclusion chromatography to remove unencapsulated dye. The pH responsiveness of FITC was validated over the relevant pH range before transport measurements. For proton-gradient measurements, liposomes were prepared with an internal pH of 5 and exposed to external buffer at pH 7. FITC fluorescence was monitored over time to detect changes in intraliposomal pH. Wild-type TMEM165, E108G TMEM165, and protein-free liposomes were compared under the same conditions. An increase in FITC fluorescence was interpreted as an increase in internal pH, consistent with proton movement out of the liposome lumen.

### Electrochemical impedance spectroscopy

Electrochemical impedance spectroscopy was performed using an Autolab PGSTAT302N potentiostat. POPC SLBs were formed on PEDOT:PSS-coated gold electrodes, and GDT1 or TMEM165 was synthesized directly into the supported bilayer using CFPS. Impedance spectra were collected before SLB formation, after SLB formation, after cell-free protein synthesis, and after ion addition. The applied sinusoidal voltage was swept from 10⁶ Hz to 1 Hz to monitor changes in the electrical response of the supported membrane. Before SLB formation, the PEDOT:PSS electrode response was fit with an RC circuit. After bilayer formation, spectra were fit with an RC(RC) equivalent circuit in which the membrane was represented as a parallel resistor-capacitor element. Membrane resistance and capacitance were extracted from the fitted circuit. Changes in membrane resistance after ion addition were used to quantify ion-dependent responses. Percent change in membrane resistance was calculated relative to the membrane resistance before ion addition. Calcium- and cobalt-dependent responses were compared for GDT1-containing SLBs, TMEM165-containing SLBs, and no-plasmid controls.

### Structural modeling and electrostatic analysis

Structural models of wild-type TMEM165 and E108G TMEM165 were generated from AlphaFold-predicted TMEM165 structures. The E108G substitution was introduced in silico, and the structures were subjected to AMBER-based minimization. A calcium ion was placed near the putative ion-coordination region formed by conserved acidic residues, including Glu108, Asp111, Glu248, and Asp251. Membrane positioning was performed using the PPM server with a DOPC bilayer model. Structural overlays were used to compare wild-type and E108G TMEM165, with emphasis on the putative pore region and acidic coordination site. Electrostatic surface potential maps were generated for the wild-type and E108G models to compare the local charge environment surrounding the predicted ion-binding region.

### Kinetic analysis

Transport-associated fluorescence traces were analyzed by fitting the initial post-addition fluorescence increase before detergent addition to a single-exponential function:

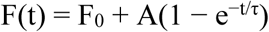

where F(t) is fluorescence at time t, F0 is the baseline fluorescence, A is the fitted amplitude, and τ is the apparent time constant. Apparent transport rates were calculated as 1/τ where indicated. These fits were used as empirical descriptions of the observed transport-associated fluorescence response rather than full mechanistic kinetic models, because concentration-dependent transport measurements were not performed. For comparisons across TMEM165 variants, apparent transport rates were normalized to relative protein expression determined by western blot densitometry. For pH-dependent experiments, steady-state fluorescence values were normalized to the maximum fluorescence observed after Triton X-100 permeabilization.

### Statistical analysis

Data are shown as mean ± standard deviation unless otherwise indicated. Fluorescence transport experiments were performed using three independent reactions unless otherwise stated. For antibody-accessibility measurements, 18 images were analyzed per condition across three independently prepared substrates. Statistical comparisons were performed using Welch’s t test. Where multiple comparisons were made, Bonferroni correction was applied. Significance was reported as ***p < 0.001, **p < 0.01, *p < 0.05, and NS, p > 0.05.

## Author contributions

Conceptualization: A.B., A.R.O., E.S., P.M., J.C., S.D

Methodology: A.B., A.R.O., Z.C., O.A., E.S., J.C., A.G., GVR

Investigation: A.B., A.R.O., Z.C., O.A., E.S.

Visualization: ES Supervision: ES, PM, SD

Writing-original draft: ES

Writing-review & editing: ES, JC, PM, SD

## Data and materials availability

All data needed to evaluate the conclusions in the paper are present in the paper and/or the Supplementary Materials. Additional data and materials are available from the corresponding authors upon reasonable request. Plasmids and other materials generated in this study are available upon reasonable request and may require a material transfer agreement.

## References

(1) Festa, M.; Minicozzi, V.; Boccaccio, A.; Lagostena, L.; Gradogna, A.; Qi, T.; Costa, A.; Larisch, N.; Hamamoto, S.; Pedrazzini, E. Current methods to unravel the functional properties of lysosomal ion channels and transporters. Cells 2022, 11 (6), 921.

(2) Li, J.; Zhang, J.; Bui, S.; Ahat, E.; Kolli, D.; Reid, W.; Xing, L.; Wang, Y. Common assays in mammalian Golgi studies. In *Golgi: Methods and Protocols*, Springer, 2022; pp 303–332.

(3) Thines, L.; Deschamps, A.; Sengottaiyan, P.; Savel, O.; Stribny, J.; Morsomme, P. The yeast protein Gdt1p transports Mn2+ ions and thereby regulates manganese homeostasis in the Golgi. Journal of Biological Chemistry 2018, 293 (21), 8048–8055.

(4) Colinet, A.-S.; Sengottaiyan, P.; Deschamps, A.; Colsoul, M.-L.; Thines, L.; Demaegd, D.; Duchêne, M.-C.; Foulquier, F.; Hols, P.; Morsomme, P. Yeast Gdt1 is a Golgi-localized calcium transporter required for stress-induced calcium signaling and protein glycosylation. Scientific reports 2016, 6 (1), 24282.

(5) Demaegd, D.; Foulquier, F.; Colinet, A.-S.; Gremillon, L.; Legrand, D.; Mariot, P.; Peiter, E.; Van Schaftingen, E.; Matthijs, G.; Morsomme, P. Newly characterized Golgi-localized family of proteins is involved in calcium and pH homeostasis in yeast and human cells. Proceedings of the National Academy of Sciences 2013, 110 (17), 6859–6864.

(6) Jankauskas, S. S.; Varzideh, F.; Kansakar, U.; Al Tibi, G.; Agyapong, E. D.; Gambardella, J.; Santulli, G. Insights into molecular and cellular functions of the Golgi calcium/manganese-proton antiporter TMEM165. Journal of Biological Chemistry 2024, 300 (8), 107567.

(7) Rosnoblet, C.; Legrand, D.; Demaegd, D.; Hacine-Gherbi, H.; de Bettignies, G.; Bammens, R.; Borrego, C.; Duvet, S.; Morsomme, P.; Matthijs, G. Impact of disease-causing mutations on TMEM165 subcellular localization, a recently identified protein involved in CDG-II. Human molecular genetics 2013, 22 (14), 2914–2928.

(8) Potelle, S.; Morelle, W.; Dulary, E.; Duvet, S.; Vicogne, D.; Spriet, C.; Krzewinski-Recchi, M.-A.; Morsomme, P.; Jaeken, J.; Matthijs, G. Glycosylation abnormalities in Gdt1p/TMEM165 deficient cells result from a defect in Golgi manganese homeostasis. Human molecular genetics 2016, 25 (8), 1489–1500.

(9) Foulquier, F.; Amyere, M.; Jaeken, J.; Zeevaert, R.; Schollen, E.; Race, V.; Bammens, R.; Morelle, W.; Rosnoblet, C.; Legrand, D. TMEM165 deficiency causes a congenital disorder of glycosylation. The American Journal of Human Genetics 2012, 91 (1), 15–26.

(10) Legrand, D.; Herbaut, M.; Durin, Z.; Brysbaert, G.; Bardor, M.; Lensink, M. F.; Foulquier, F. New insights into the pathogenicity of TMEM165 variants using structural modeling based on AlphaFold 2 predictions. Computational and Structural Biotechnology Journal 2023, 21, 3424–3436.

(11) Motta, I.; Gohlke, A.; Adrien, V.; Li, F.; Gardavot, H.; Rothman, J. E.; Pincet, F. Formation of giant unilamellar proteo-liposomes by osmotic shock. Langmuir 2015, 31 (25), 7091–7099.

(12) Diederichs, T.; Tampé, R. Single Cell-like systems reveal active unidirectional and Light-Controlled transport by nanomachineries. ACS nano 2021, 15 (4), 6747–6755.

(13) Tosaka, T.; Kamiya, K. Function investigations and applications of membrane proteins on artificial lipid membranes. International journal of molecular sciences 2023, 24 (8), 7231.

(14) Selivanovitch, E.; Ostwalt, A.; Chao, Z.; Daniel, S. Emerging Designs and Applications for Biomembrane Biosensors. Annual Review of Analytical Chemistry 2024, 17 (1), 339–366.

(15) Dos Santos, E. C.; Belluati, A.; Necula, D.; Scherrer, D.; Meyer, C. E.; Wehr, R. P.; Lörtscher, E.; Palivan, C. G.; Meier, W. Combinatorial Strategy for Studying Biochemical Pathways in Double Emulsion Templated Cell-Sized Compartments. Advanced Materials 2020, 32 (48), 2004804.

(16) Dam, T.; Chouliara, M.; Junghans, V.; Jönsson, P. Supported lipid bilayers and the study of two-dimensional binding kinetics. Frontiers in molecular biosciences 2022, 9, 833123.

(17) Liu, H.-Y.; Pappa, A.-M.; Pavia, A.; Pitsalidis, C.; Thiburce, Q.; Salleo, A.; Owens, R. M.; Daniel, S. Self-assembly of mammalian-cell membranes on bioelectronic devices with functional transmembrane proteins. Langmuir 2020, 36 (26), 7325–7331.

(18) Pappa, A.-M.; Liu, H.-Y.; Traberg-Christensen, W.; Thiburce, Q.; Savva, A.; Pavia, A.; Salleo, A.; Daniel, S.; Owens, R. M. Optical and electronic ion channel monitoring from native human membranes. ACS nano 2020, 14 (10), 12538–12545.

(19) Richards, M. J.; Hsia, C.-Y.; Singh, R. R.; Haider, H.; Kumpf, J.; Kawate, T.; Daniel, S. Membrane protein mobility and orientation preserved in supported bilayers created directly from cell plasma membrane blebs. Langmuir 2016, 32 (12), 2963–2974.

(20) Shimizu, Y.; Inoue, A.; Tomari, Y.; Suzuki, T.; Yokogawa, T.; Nishikawa, K.; Ueda, T. Cell-free translation reconstituted with purified components. Nature biotechnology 2001, 19 (8), 751–755.

(21) Kuruma, Y.; Ueda, T. The PURE system for the cell-free synthesis of membrane proteins. Nature protocols 2015, 10 (9), 1328–1344.

(22) Harris, N. J.; Booth, P. J. Investigating the Insertion and Folding of Membrane Transporters into Lipid Bilayers Using a Cell Free Expression System. Biophysical Journal 2017, 112 (3), 204a.

(23) Harris, N. J.; Pellowe, G. A.; Booth, P. J. Cell-free expression tools to study co-translational folding of alpha helical membrane transporters. Scientific reports 2020, 10 (1), 9125.

(24) Manzer, Z. A.; Ghosh, S.; Jacobs, M. L.; Krishnan, S.; Zipfel, W. R.; Piñeros, M.; Kamat, N. P.; Daniel, S. Cell-free synthesis of a transmembrane mechanosensitive channel protein into a hybrid-supported lipid bilayer. ACS Applied Bio Materials 2021, 4 (4), 3101–3112.

(25) Manzer, Z. A.; Ghosh, S.; Roy, A.; Jacobs, M. L.; Carten, J.; Kamat, N. P.; Daniel, S. Cell-free synthesis goes electric: dual optical and electronic biosensor via direct channel integration into a supported membrane electrode. ACS synthetic biology 2023, 12 (2), 502–510.

(26) Harris, N. J.; Reading, E.; Booth, P. J. Cell-free synthesis strategies to probe co-translational folding of proteins within lipid membranes. In *Cell-Free Gene Expression: Methods and Protocols*, Springer, 2022; pp 273–292.

(27) Lu, Z.; van Niekerk, D.; Savva, A.; Kallitsis, K.; Thiburce, Q.; Salleo, A.; Pappa, A.-M.; Owens, R. M. Understanding electrochemical properties of supported lipid bilayers interfaced with organic electronic devices. Journal of Materials Chemistry C 2022, 10 (20), 8050–8060.

(28) Atanasov, V.; Knorr, N.; Duran, R. S.; Ingebrandt, S.; Offenhäusser, A.; Knoll, W.; Köper, I. Membrane on a chip: A functional tethered lipid bilayer membrane on silicon oxide surfaces. Biophysical journal 2005, 89 (3), 1780–1788.

(29) Lebredonchel, E.; Houdou, M.; Potelle, S.; de Bettignies, G.; Schulz, C.; Recchi, M.-A. K.; Lupashin, V.; Legrand, D.; Klein, A.; Foulquier, F. Dissection of TMEM165 function in Golgi glycosylation and its Mn2+ sensitivity. Biochimie 2019, 165, 123–130.

(30) Stribny, J.; Thines, L.; Deschamps, A.; Goffin, P.; Morsomme, P. The human Golgi protein TMEM165 transports calcium and manganese in yeast and bacterial cells. Journal of Biological Chemistry 2020, 295 (12), 3865–3874.

(31) Bastin, G.; Singh, K.; Dissanayake, K.; Mighiu, A. S.; Nurmohamed, A.; Heximer, S. P. Amino-terminal cysteine residues differentially influence RGS4 protein plasma membrane targeting, intracellular trafficking, and function. Journal of Biological Chemistry 2012, 287 (34), 28966–28974.

(32) Li, G.-W.; Oh, E.; Weissman, J. S. The anti-Shine–Dalgarno sequence drives translational pausing and codon choice in bacteria. Nature 2012, 484 (7395), 538–541.

(33) Ikemura, T. Correlation between the abundance of Escherichia coli transfer RNAs and the occurrence of the respective codons in its protein genes. Journal of molecular biology 1981, 146 (1), 1–21.

(34) Thines, L.; Stribny, J.; Morsomme, P. From the uncharacterized protein family 0016 to the GDT1 family: molecular insights into a newly-characterized family of cation secondary transporters. Microbial Cell 2020, 7 (8), 202.

(35) Deschamps, A.; Thines, L.; Colinet, A.-S.; Stribny, J.; Morsomme, P. The yeast Gdt1 protein mediates the exchange of H+ for Ca2+ and Mn2+ influencing the Golgi pH. Journal of Biological Chemistry 2023, 299 (5).

(36) Zajac, M.; Mukherjee, S.; Anees, P.; Oettinger, D.; Henn, K.; Srikumar, J.; Zou, J.; Saminathan, A.; Krishnan, Y. A mechanism of lysosomal calcium entry. Science Advances 2024, 10 (7), eadk2317.

(37) Chen, R.; Liu, B.; Jaślan, D.; Kucej, L.; Kudrina, V.; Warnke, B.; Klingl, Y. E.; Petrauskas, A.; Castro, S. P.; Maeda, K. Lysosomal TMEM165 controls cellular ion homeostasis and survival by mediating lysosomal Ca2+ import and H+ efflux. Nature Communications 2025, 16 (1), 5209.

