## Supplemental File for "Cell-free membrane systems reveal electrostatic control of ion transport in TMEM165 and disease-associated CaCA2 variants"

**Table S1.** DNA sequences for pET28a-6xHis-<sup>wt</sup>TMEM165-TEV-GFP.

| Construct | DNA Sequence |
| --- | --- |
| pET28a-6xHis-<sup>wt</sup>TMEM165-TEV-GFP | ATGGGCAGCAGCCATCATCATCATCATCACAGCAGCGGCCTGGTGCCGC<br>GCGGCAGCCATATGGCTAGCATGACTGGTGGACAGCAAATGGGTCGCGG<br>ATCCATGGCCGCAGCGGCCCCTGGTAACGGTCGGGCCTCAGCCCCACGC<br>CTGTTGCTGCTGTTCTTAGTCCCGCTGCTGTGGGCCCCAGCCGCCGTGC<br>GTGCAGGCCCGGATGAAGATCTGAGTCACAGAAATAAAGAACCACCTG<br>CACCTGCACAGCAATTACAGCCGCAGCCGGTTGCTGTTCAGGGTCCTGA<br>ACCGGCTCGTGTTGAAAAAATATTTACACCTGCAGCTCCAGTACACACA<br>AATAAAGAGGATCCTGCAACGCAGACAAATCTGGGTTTTATTCATGCGTT<br>TGTTGCCGCAATATCCGTTATTATTGTATCAGAGTTGGGTGATAAAACATT<br>CTTCATCGCCGCGATCATGGCTATGCGTTATAATCGTCTTACGGTCCTGGC<br>CGGTGCGATGCTGGCTTTAGGTCTGATGACCTGTCTGTCTGTTCTCTTCG<br>GCTATGCCACCACCGTGATCCCGCGTGTGTATACTTATTACGTCAGTACA<br>GTGCTTTTCGCGATCTTCGGTATACGGATGCTGAGAGAAGGACTTAAGAT<br>GAGCCCTGACGAAGGTCAAGAAGAATTAGAGGAAGTACAGGCGGAACT<br>GAAGAAAAAAGATGAAGAATTTCAACGCACCAAATTACTGAATGGTCC<br>AGGAGACGTTGAGACGGGAACTTCTATCACCGTACCTCAAAAAAAATG<br>GCTGCATTTCATTAGCCCTATCTTTGTCCAAGCCTTGACACTCACGTTTCT<br>TGCGGAATGGGGCGATCGCTCCCAGTTAACCACAATTGTACTGGCGGCT<br>CGCGAAGATCCGTATGGCGTTGCGGTGGGCGGCACCGTTGGACACTGTC<br>TGTGTACAGGTCTCGCTGTAATCGGCGGTAGAATGATTGCGCAAAAAAT<br>ATCAGTCCGCACGGTAACTATAATTGGCGGCATTGTCTTTCTTGCCTTTG<br>CTTTTTCTGCTCTGTTTATCTCTCCAGATTCCGGTTTCGGGTCTGGTGGTG<br>GCTCCGAGAACTTATACTTTCAGGGTATGAGCAAAGGGGAAGAGTTGTT<br>TACGGGTGTTGTACCCATCTTGGTAGAGTTAGATGGGGATGTGAATGGAC<br>ACAAATTCTCGGTCCGCGGAGAGGGTGAAGGGGATGCTACTAATGGTAA<br>ATTAACACTGAAATTCATTTGTACTACAGGTAAATTGCCGGTTCCGTGGC<br>CGACATTAGTTACCACTCTCACGTACGGTGTTCAATGTTTTTCACGTTAC<br>CCTGACCACATGAAACGGCACGATTTTTTTAAGTCCGCTATGCCTGAGG<br>GATATGTTCAGGAGCGGACAATTAGCTTCAAAGATGATGGCACGTATAA<br>AACACGCGCAGAAGTTAAGTTCGAAGGGGATACCCTCGTTAATCGTATC<br>GAATTAAAAGGGATAGATTTCAAAGAGGACGGGAACATCTTAGGGCATC<br>TGGAATATAACTTTAACTCTCATAATGTATACATTACAGCTGACAAACAA<br>AAAAATGGTATAAAAGCAAACTTTAAAATTAGACACAATGTCGAAGACG<br>GCTCTGTCCAGTTGGCAGATCACTATCAGCAGAATACACCTATCGGTGAC<br>GGACCGGTTCTTCTGCCTGACAATCATTATCTGTCAACGCAGTCTGTTTT<br>ATCGAAAGACCCTAATGAAAAACGAGATCACATGGTCCTGCTTGAATTC<br>GTCACCGCAGCAGGCATAACACACGGAATGGACGAATTATACAAATGAC<br>TCGAGCACCACCACCACCACCACTGA |

**Table S2.** DNA sequences for Flag-<sup>wt</sup>TMEM165-6xHis

| Construct | DNA Sequence |
| --- | --- |
| Flag-<sup>wt</sup>TMEM165-6xHis | ATGGACTATAAGGACGATGATGATAAAGCAGCAGCAGCCCCT<br>GGTAATGGTCGTGCAAGCGCACCGCGTCTGCTGCTGTTATTT<br>CTGGTTCCGCTGCTGTGGGCACCAGCAGCAGTTCGTGCAGG<br>TCCGGATGAAGATCTGAGCCATCGTAATAAAGAACCGCCTGC<br>ACCGGCACAGCAGCTGCAGCCGCAGCCGGTTGCAGTTCAGG<br>GTCCTGAACCGGCACGTGTTGAAAAGATTTTTACACCGGCA<br>GCACCGGTTCATACCAATAAAGAAGATCCGGCAACACAGAC |
|  | CAATCTGGGTTTTATTCATGCATTTGTTGCAGCCATTAGCGTG<br>ATTATTGTTAGCGAACTGGGTGACAAAACCTTTTTCATTGCA<br>GCAATTATGGCCATGCGTTATAATCGTCTGACCGTTCTGGCAG<br>GCGCAATGCTGGCACTGGGTCTGATGACCTGCCTGAGCGTTC<br>TGTTTGGTTATGCAACCACCGTTATTCCGCGTGTGTATACCTA<br>TTATGTTAGCACCGTGCTGTTTGCCATTTTTGGTATTCGTATGC<br>TGCGTGAAGGTCTGAAAATGAGCCCTGATGAAGGTCAAGAA<br>GAACTGGAAGAAGTTCAGGCAGAACTGAAAAAGAAAGATG<br>AAGAATTTCAGCGCACCAAACTGCTGAATGGTCCGGGTGAT<br>GTTGAAACCGGTACAAGCATTACCGTTCCGCAGAAAAAGTG<br>GCTGCATTTTATCAGCCCGATTTTTGTTCAGGCACTGACCCTG<br>ACATTTCTGGCAGAATGGGGTGATCGTAGCCAGCTGACCACC<br>ATTGTGCTGGCAGCACGTGAAGATCCGTATGGTGTTGCAGTT<br>GGTGGCACCGTTGGTCATTGTCTGTGTACCGGTCTGGCAGTT<br>ATTGGTGGTCGTATGATTGCACAGAAAATTAGCGTTCGTACC<br>GTGACCATTATTGGCGGTATTGTTTTTCTGGCATTTGCATTTAG<br>CGCACTGTTTATTAGTCCGGATAGCGGTTTTGGTGGTAGCCAT<br>CACCATCATCATCATTAA |

**Table S3.** DNA sequences for clinically relevant mutants of Flag-<sup>wt</sup>TMEM165-6xHis .

| Construct | DNA Sequence |
| --- | --- |
| Patients 1-3<br>(c.792+182<br>G>A) | ATGGACTATAAGGACGATGATGATAAAGCAGCAGCAGCCCCTGGTAATGGTC<br>GTGCAAGCGCACCGCGTCTGCTGCTGTTATTTCTGGTTCCGCTGCTGTGGGC<br>ACCAGCAGCAGTTCGTGCAGGTCCGGATGAAGATCTGAGCCATCGTAATAA<br>AGAACCGCCTGCACCGGCACAGCAGCTGCAGCCGCAGCCGGTTGCAGTTC<br>AGGGTCCTGAACCGGCACGTGTTGAAAAGATTTTTACACCGGCAGCACCGG<br>TTCATACCAATAAAGAAGATCCGGCAACACAGACCAATCTGGGTTTTATTCA<br>TGCATTTGTTGCAGCCATTAGCGTGATTATTGTTAGCGAACTGGGTGACAAA<br>ACCTTTTTCATTGCAGCAATTATGGCCATGCGTTATAATCGTCTGACCGTTCT<br>GGCAGGCGCAATGCTGGCACTGGGTCTGATGACCTGCCTGAGCGTTCTGTT<br>TGGTTATGCAACCACCGTTATTCCGCGTGTGTATACCTATTATGTTAGCACCG<br>TGCTGTTTGCCATTTTTGGTATTTCGTATGCTGCGTGAAGGTCTGAAAATGAG<br>CCCTGATGAAGGTCAAGAAGAAGTTCAGGCAGAACTGAAAA<br>AGAAAGACGAAGAGAAAACCCTGGATACCCTGAAATGGGCAACCGCATTTC<br>TGCACATGCAGGATTGTTTTACCAGCGATAATAGCGCAGCATTAGGTGGTAG<br>CCATCACCATCATCATCATTAA |
| Patient 4<br>(<sup>126</sup>Arg>His) | ATGGACTATAAGGACGATGATGATAAAGCAGCAGCAGCCCCTGGTAATGGTC<br>GTGCAAGCGCACCGCGTCTGCTGCTGTTATTTCTGGTTCCGCTGCTGTGGGC<br>ACCAGCAGCAGTTCGTGCAGGTCCGGATGAAGATCTGAGCCATCGTAATAA<br>AGAACCGCCTGCACCGGCACAGCAGCTGCAGCCGCAGCCGGTTGCAGTTC<br>AGGGTCCTGAACCGGCACGTGTTGAAAAGATTTTTACACCGGCAGCACCGG<br>TTCATACCAATAAAGAAGATCCGGCAACACAGACCAATCTGGGTTTTATTCA<br>TGCATTTGTTGCAGCCATTAGCGTGATTATTGTTAGCGAACTGGGTGACAAA<br>ACCTTTTTCATTGCAGCAATTATGGCCATGCGCTATAATCATCTGACCGTTCT<br>GGCAGGCGCAATGCTGGCACTGGGTCTGATGACCTGTCTGAGCGTTCTGTT<br>TGGTTATGCAACCACCGTTATTCCGCGTGTGTATACCTATTATGTTAGCACCG<br>TGCTGTTTGCCATTTTTGGTATTCGTATGCTGCGTGAAGGTCTGAAAATGAG |
|  | CCCTGATGAAGGTCAAGAAGAAGTTCAGGCAGAACTGAAAA<br>AGAAAGATGAAGAATTTTCAGCGCACCAAAGTCTGAATGGTCCGGGTGATG<br>TTGAAACCGGTACAAGCATTACCGTTCCGCAGAAAAAGTGGCTGCATTTTAT<br>CAGCCCGATTTTGTTCAGGCACTGACCCTGACATTTCTGGCAGAATGGGGT<br>GATCGTAGCCAGCTGACCACCATTTGTGCTGGCAGCACGTGAAGATCCGTATG<br>GTGTTGCAGTTGGTGGCACCGTTGGTCATTGTCTGTGTACCGGTCTGGCAGT<br>TATTGGTGGTCGTATGATTGCACAGAAAATTAGCGTTCGTACCGTGACCATTA<br>TTGGCGGTATTGTTTTTCTGGCATTTCATTAGCGCACTGTTTATTAGTCCG<br>GATAGCGGTTTTGGTGGTAGCCATCACCATCATCATCATTA |
| Patient 5<br>(<sup>304</sup>Gly>Arg) | ATGGACTATAAGGACGATGATGATAAAGCAGCAGCAGCCCCTGGTAATGGTC<br>GTGCAAGCGCACCGCGTCTGCTGCTGTTATTTCTGGTTCCGCTGCTGTGGGC<br>ACCAGCAGCAGTTTCGTGCAGGTCCGGATGAAGATCTGAGCCATCGTAATAA<br>AGAACCGCCTGCACCGGCACAGCAGCTGCAGCCGCAGCCGTTGCAGTTC<br>AGGGTCCTGAACCGGCACGTGTTGAAAAGATTTTACACCGGCAGCACCGG<br>TTCATACCAATAAAGAAGATCCGGCAACACAGACCAATCTGGGTTTTATTCA<br>TGCATTTGTTGCAGCCATTAGCGTGATTATTGTTAGCGAACTGGGTGACAAA<br>ACCTTTTTTCATTGCAGCAATTATGGCCATGCGTTATAATCGTCTGACCGTTCT<br>GGCAGGCGCAATGCTGGCACTGGGTCTGATGACCTGCCTGAGCGTTCTGTT<br>TGTTTATGCAACCACCGTTATTCCGCGTGTGTATACCTATTATGTTAGCACCG<br>TGCTGTTTGCCATTTTTGGTATTTCGTATGCTGCGTGAAGGTCTGAAAATGAG<br>CCCTGATGAAGGTCAAGAAGAAGTTCAGGCAGAACTG<br>AAAAAGAAAGATGAAGAATTTTCAGCGCACCAAAGTCTGAATGGTCCGGG<br>TGATGTTGAAACCGGTACAAGCATTACCGTTCCGCAGAAAAAGTGGCTGCA<br>TTTTATCAGCCCGATTTTTGTTCAGGCACTGACCCTGACATTTCTGGCAGAAT<br>GGGGTGATCGTAGCCAGCTGACCACCATTTGTGCTGGCAGCACGTGAAGATC<br>CGTATGGTGTGTCAGTTGGTGGCACCGTTGGTCATTGTCTGTGTACCGGTCT<br>GGCAGTTATTGGTGGTCGTATGATTGCACAGAAAATTAGCGTTCGTACCGTG<br>ACCATTATTCGCGGTATTGTTTTTCTGGCATTTCATTAGCGCACTGTTTATT<br>AGTCCGGATAGCGGTTTTGGTGGTAGCCATCACCATCATCATCATTA |
| Patient 5<br>(<sup>126</sup>Arg>Cys) | ATGGACTATAAGGACGATGATGATAAAGCAGCAGCAGCCCCTGGTAATGGTC<br>GTGCAAGCGCACCGCGTCTGCTGCTGTTATTTCTGGTTCCGCTGCTGTGGGC<br>ACCAGCAGCAGTTCGTGCAGGTCCGGATGAAGATCTGAGCCATCGTAATAA<br>AGAACCGCCTGCACCGGCACAGCAGCTGCAGCCGCAGCCGGTTGCAGTTC<br>AGGGTCCTGAACCGGCACGTGTTGAAAAGATTTTTACACCGGCAGCACCGG<br>TTCATACCAATAAAGAAGATCCGGCAACACAGACCAATCTGGGTTTTATTCA<br>TGCATTTGTTGCAGCCATTAGCGTGATTATTGTTAGCGAACTGGGTGACAAA<br>ACCTTTTTCATTGCAGCAATTATGGCCATGCGTTATAATTGTCTGACCGTTCT<br>GGCAGGCGCAATGCTGGCACTGGGTCTGATGACCTGCCTGAGCGTTCTGTT<br>TGGTTATGCAACCACCGTTATTCCGCGTGTGTATACCTATTATGTTAGCACCG<br>TGCTGTTTGCCATTTTTGGTATTTCGTATGCTGCGTGAAGGTCTGAAAATGAG<br>CCCTGATGAAGGTCAAGAAGAAGTTCAGGCAGAACTGAAAA<br>AGAAAGATGAAGAATTTTCAGCGCACCAAAGTCTGAATGGTCCGGGTGATG<br>TTGAAACCGGTACAAGCATTACCGTTCCGCAGAAAAAGTGGCTGCATTTTAT<br>CAGCCCGATTTTGTTCAGGCACTGACCCTGACATTTCTGGCAGAATGGGGT<br>GATCGTAGCCAGCTGACCACCATTTGTGCTGGCAGCACGTGAAGATCCGTATG<br>GTGTTGCAGTTGGTGGCACCGTTGGTCATTGTCTGTGTACCGGTCTGGCAGT<br>TATTGGTGGTCGTATGATTGCACAGAAAATTAGCGTTCGTACCGTGACCATTA<br>TTGGCGGTATTGTTTTTCTGGCATTTCATTAGCGCACTGTTTATTAGTCCG<br>GATAGCGGTTTTGGTGGTAGCCATCACCATCATCATCATTA |
| Patient 6<br>(<sup>108</sup>Glu>Gly) | ATGGACTATAAGGACGATGATGATAAAGCAGCAGCAGCCCCTGGTAATGGTC<br>GTGCAAGCGCACCGCGTCTGCTGCTGTTATTTCTGGTTCCGCTGCTGTGGGC |
|  | ACCAGCAGCAGTTCGTGCAGGTCCGGATGAAGATCTGAGCCATCGTAATAA<br>AGAACCGCCTGCACCGGCACAGCAGCTGCAGCCGCAGCCGGTTGCAGTTC<br>AGGGTCCTGAACCGGCACGTGTTGAAAAGATTTTTACACCGGCAGCACCGG<br>TTCATACCAATAAAGAAGATCCGGCAACACAGACCAATCTGGGTTTTATTCA<br>TGCATTTGTTGCAGCCATTAGCGTGATTATTGTTAGCGGTCTGGGTGATAAAA<br>CCTTTTTCATTGCAGCAATTATGGCCATGCGTTATAATCGTCTGACCGTTCTG<br>GCAGGCGCAATGCTGGCACTGGGTCTGATGACCTGCCTGAGCGTTCTGTTT<br>GGTTATGCAACCACCGTTATTCCGCGTGTGTATACCTATTATGTTAGCACCGT<br>GCTGTTTGCCATTTTTGGTATTCGTATGCTGCGTGAAGGTCTGAAAATGAGC<br>CCTGATGAAGGTCAAGAAGAACTGGAAGAAGTTCAGGCAGAACTGAAAAA<br>GAAAGATGAAGAATTTCAGCGCACCAAACTGCTGAATGGTCCGGGTGATGT<br>TGAAACCGGTACAAGCATTACCGTTCCGCAGAAAAAGTGGCTGCATTTTATC<br>AGCCCGATTTTTGTTCAGGCACTGACCCTGACATTTCTGGCAGAATGGGGTG<br>ATCGTAGCCAGCTGACCACCATTGTGCTGGCAGCACGTGAAGATCCGTATGG<br>TGTTGCAGTTGGTGGCACCGTTGGTCATTGTCTGTGTACCGGTCTGGCAGTT<br>ATTGGTGGTCGTATGATTGCACAGAAAATTAGCGTTCGTACCGTGACCATTAT<br>TGGCGGTATTGTTTTTCTGGCATTTGCATTTAGCGCACTGTTTATTAGTCCGG<br>ATAGCGGTTTTGGTGGTAGCCATCACCATCATCATCATTAA |

**Table S4.** DNA sequences for 6xHis loading control pQE30-6xHis-IF1 (MW 9.648.06 Da)

| Construct | DNA Sequence |
| --- | --- |
| pQE30-6xHis-IF1 | ATGAGAGGATCGCATCACCATCACCATCACGGATCCATGGCCAAAGAAGACAATATT<br>GAAATGCAAGGTACCGTTCTTGAAACGTTGCCTAATACCATGTTCCGCGTAGAGTTA<br>GAAAACGGTCACGTGGTTACTGCACACATCTCCGGTAAAATGCGCAAAAACTACAT<br>CCGCATCCTGACGGGCGACAAAGTGACTGTTGAACTGACCCCGTACGACCTGAGCA<br>AAGGCCGCATTGTCTTCCGTAGTCGCTGA |

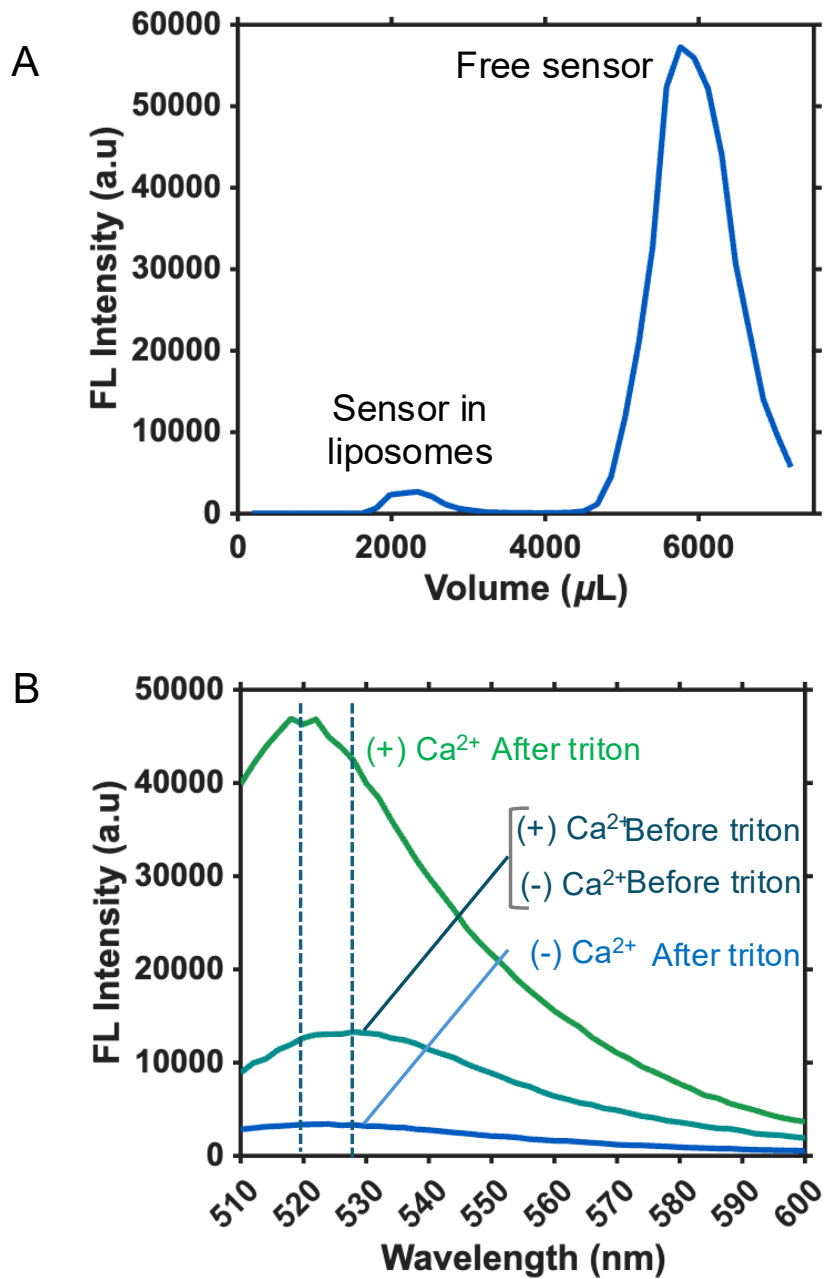

**Fig. S1. Purification and validation of cargo-loaded liposomes.** (A) Size-exclusion chromatography (SEC) was used to separate liposomes containing encapsulated fluorescent sensors from unencapsulated free sensor. The chromatogram shows the purification profile used to collect the liposome-containing fraction. (B) Emission spectra of encapsulated dye under $\pm\text{Ca}^{2+}$ and $\pm\text{Triton X-100}$ conditions were used to assess dye encapsulation and membrane integrity. Minimal fluorescence change in the absence of Triton indicates limited sensor leakage, while Triton-mediated liposome disruption reveals the total encapsulated sensor signal.

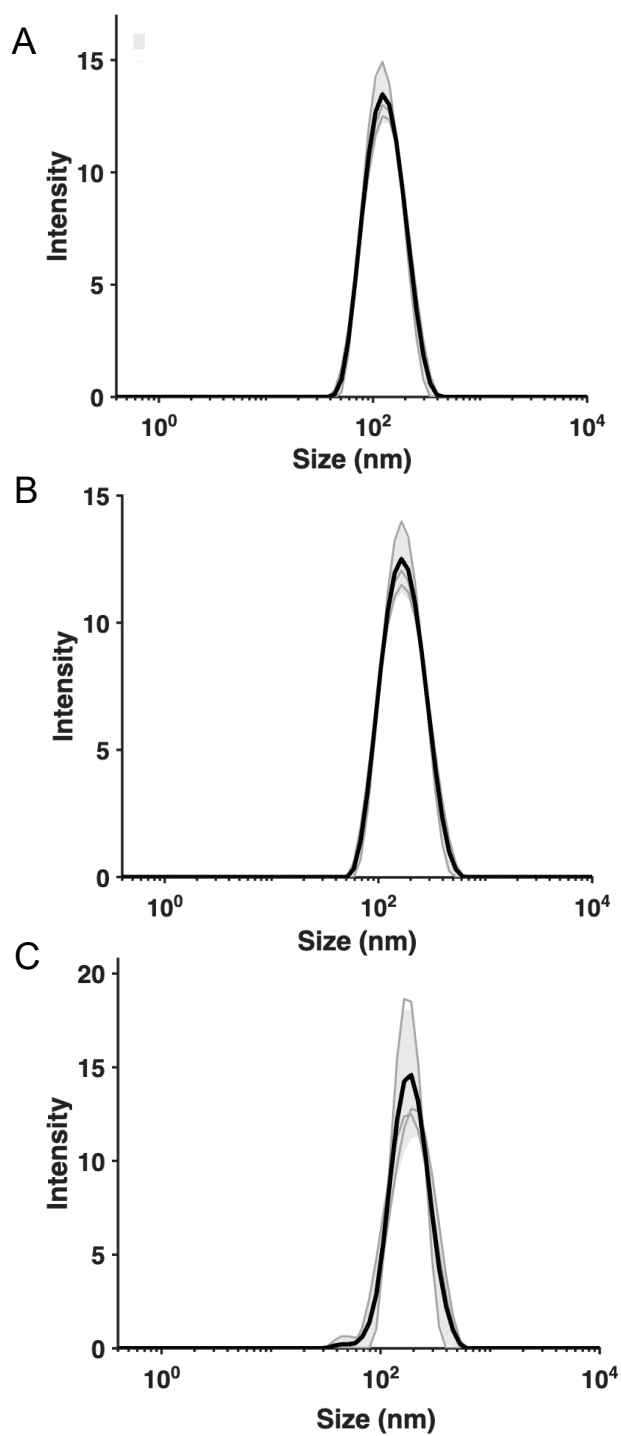

**Fig. S2. DLS profiles of liposome formulations.** (A) Liposomes prepared for supported lipid bilayer formation displayed an average diameter of approximately 100 nm. (B) Liposomes prepared for cell-free protein synthesis displayed an average diameter of approximately 200 nm. (C) Protein-containing liposomes, referred to as proteoliposomes, showed a modest increase in average diameter to approximately 220 nm following SEC purification. For each sample, three replicate DLS traces are shown in gray, with the average trace shown in black.

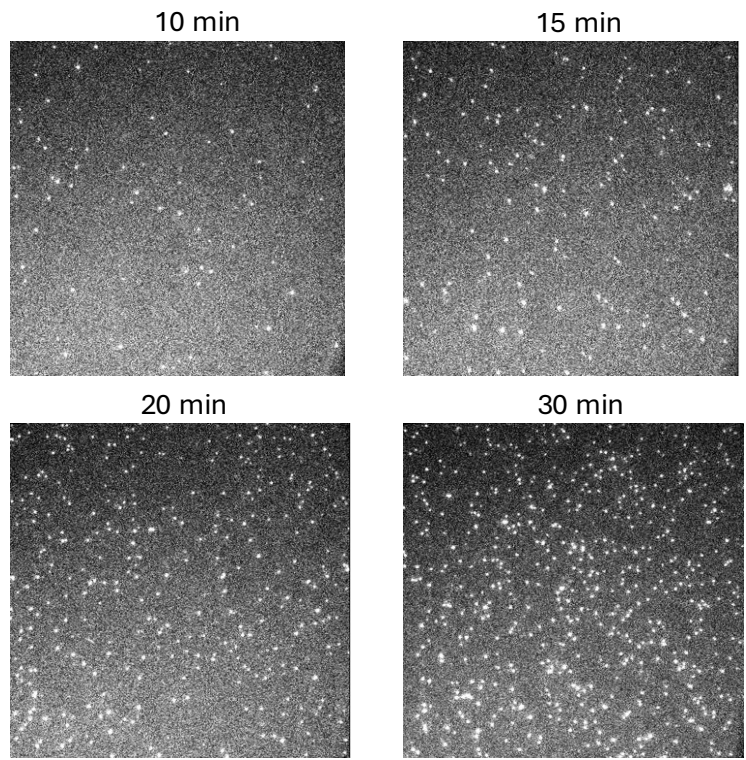

**Fig. S3. TIRF analysis of GDT1-GFP incorporation into SLBs.** TIRF microscopy was used to monitor GDT1-GFP synthesis and incorporation into SLBs over time. GDT1-GFP puncta increased with reaction time, with 20-30 min producing discrete puncta that could be readily detected and quantified with minimal background signal. Based on this analysis, a 20-30 min synthesis window was used for subsequent SLB incorporation experiments.

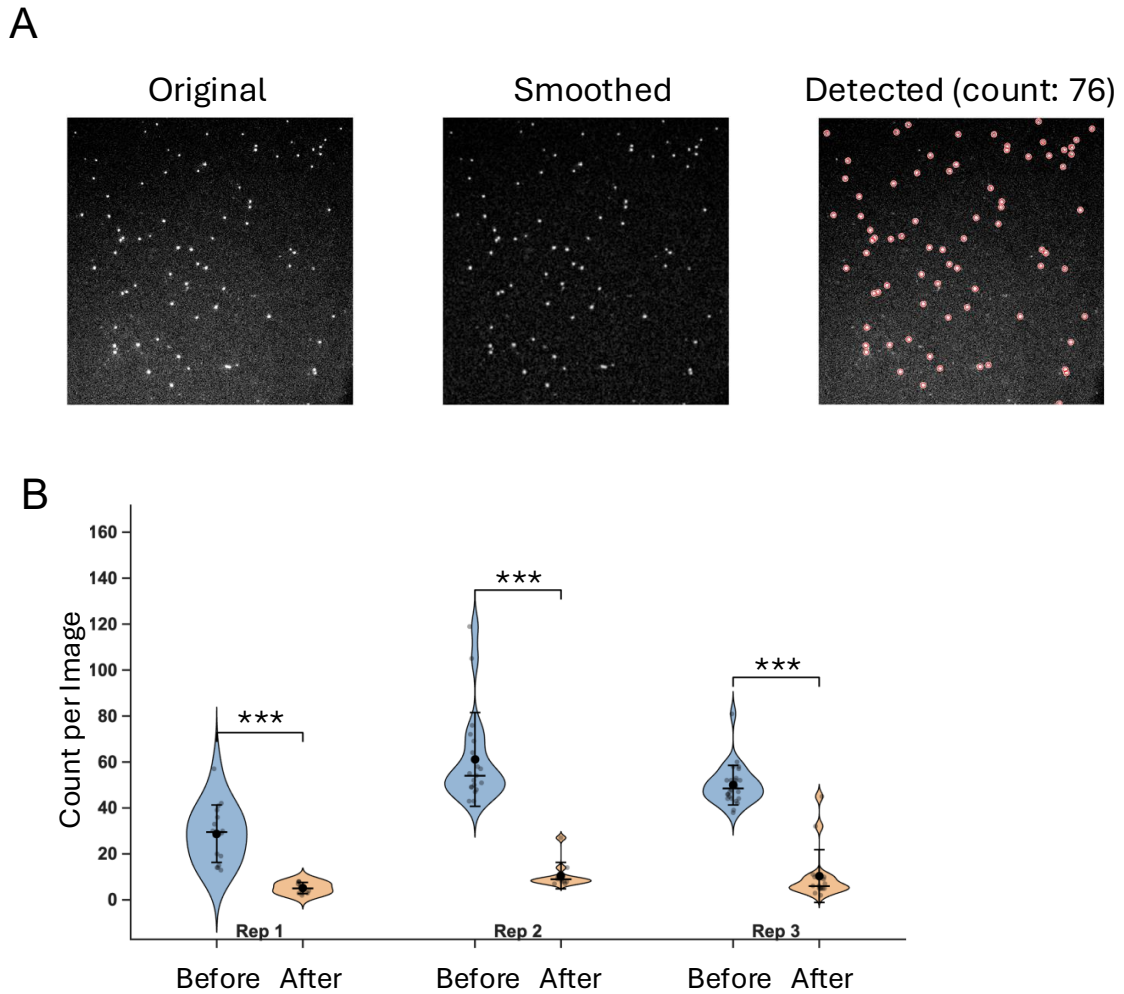

**Fig. S4. TIRF analysis of TMEM165-GFP incorporation into SLBs.** TMEM165-GFP was synthesized for 30 min prior to TIRF analysis. Fewer TMEM165-GFP puncta were observed compared with GDT1-GFP, consistent with the lower relative synthesis yield observed in the time-resolved fluorescence measurements shown in Fig. 2F–G. **(A)** Representative TIRF images showing the raw image, smoothed image, and detected particles used for particle-counting analysis. The detected puncta indicate association of TMEM165-GFP with the SLB. **(B)** To distinguish membrane-anchored TMEM165-GFP from nonspecifically adhered GFP signal, a TEV protease cleavage site was introduced between TMEM165 and GFP. Following TEV treatment, a reduction in GFP puncta was observed, supporting that a fraction of the TMEM165-GFP signal arises from membrane-anchored protein. Quantification was performed using 18 images collected from three independent biological replicates, corresponding to three independent cell-free protein synthesis reactions. Statistical significance was determined by Welch's t-test, with \*\*\* indicating $p < 0.001$.

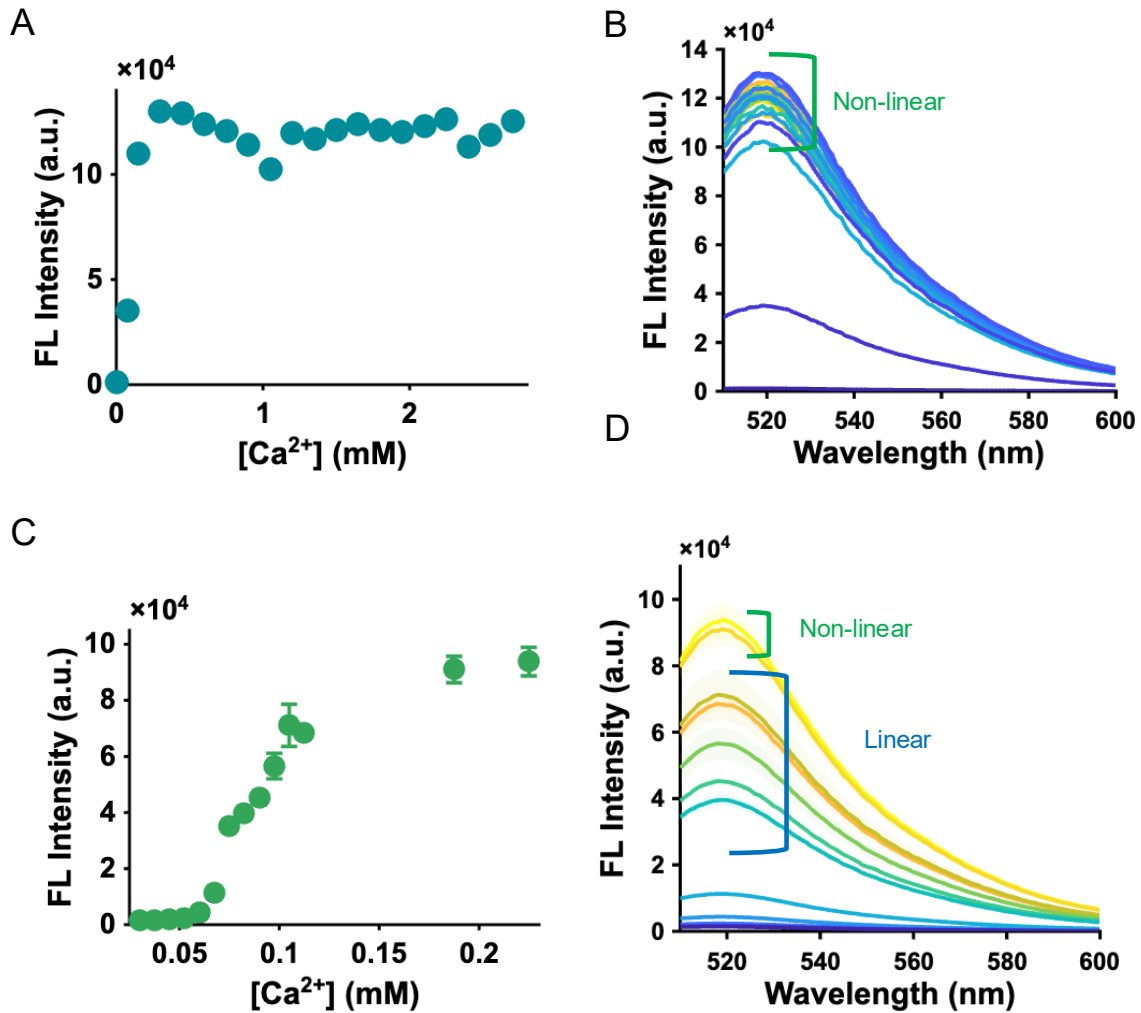

**Fig. S5. Identification of the linear response range of Fluo-4FF to calcium.** Fluo-4FF fluorescence was measured across varying $Ca^{2+}$ concentrations to identify the concentration range suitable for quantitative calcium influx measurements. **(A)** Fluorescence intensity measured at 518 nm following excitation at 464 nm showed a nonlinear response across the full $Ca^{2+}$ concentration range. **(B)** Corresponding emission spectra collected across the full $Ca^{2+}$ titration. **(C)** Fluorescence intensity plotted over the lower $Ca^{2+}$ concentration range used to identify the linear response region of the sensor. **(D)** Corresponding emission spectra within the linear response range. Measurements were performed in triplicate, and error bars represent standard deviation. Defining the linear range was necessary to ensure that subsequent fluorescence changes could be interpreted as proportional to $Ca^{2+}$ influx; outside this range, nonlinear sensor response would confound assumptions about relative transport rates.

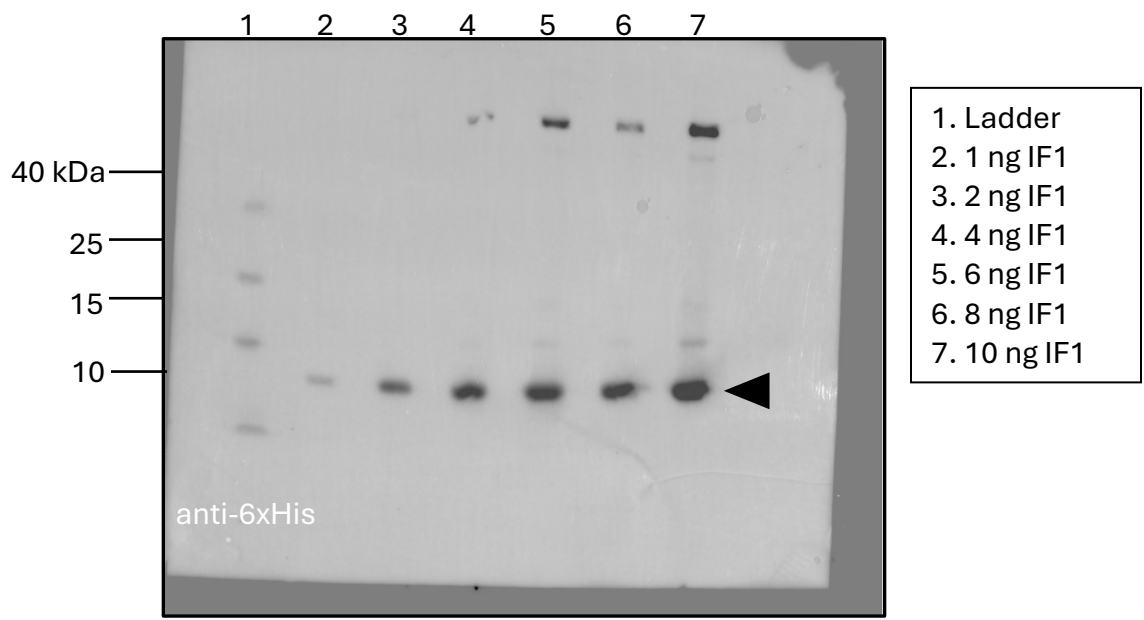

**Fig. S6. Uncropped western blot analysis of 6xHis-IF1 standard.** Uncropped western blot against the N-terminal 6xHis tag of pQE30-6xHis-IF1, abbreviated as IF1 in the figure. IF1 contains an N-terminal 6xHis tag and has an expected molecular weight of approximately 10 kDa. This blot was used to identify an appropriate IF1 standard for quantifying relative synthesis levels of TMEM165 mutants. Establishing this standard was necessary for subsequent normalization of transport activity to protein expression.

**A**

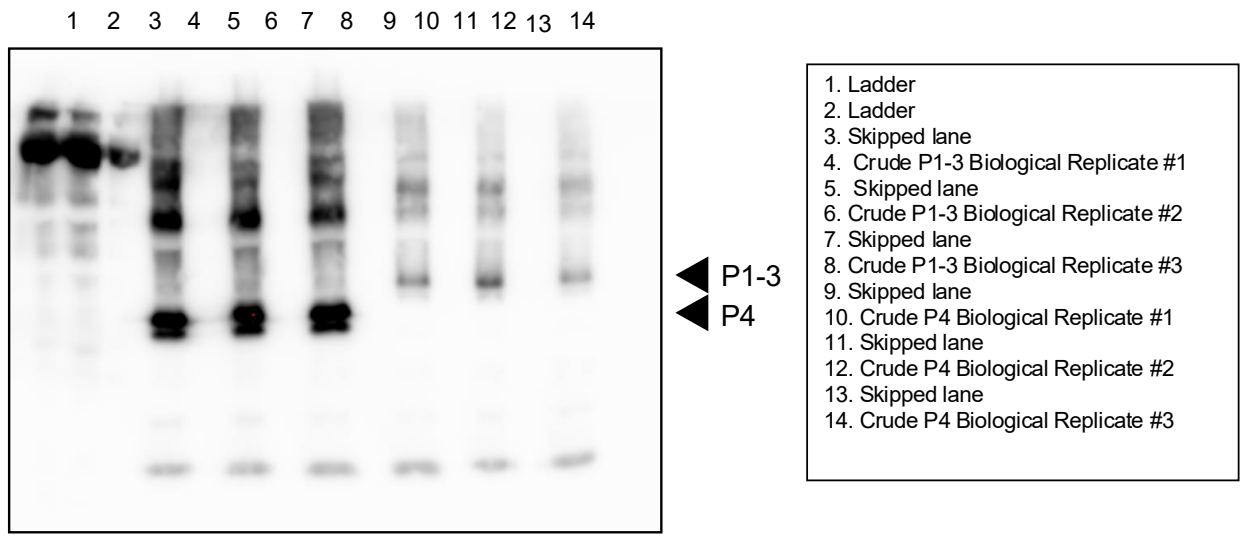

**B**

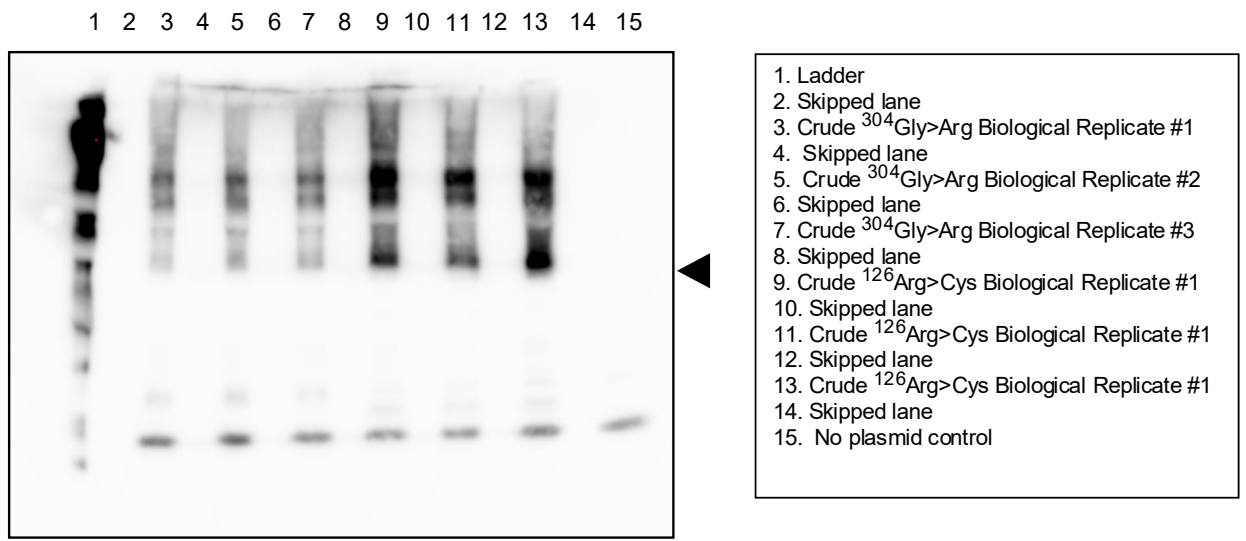

**Fig. S7. Uncropped western blots of TMEM165 variants detected by anti-His antibody.** Uncropped western blots showing anti-His detection of TMEM165 variants. **(A)** Blots corresponding to patient variants P1–P3 and P4. **(B)** Blots corresponding to the two patient 5 variants. Three independent biological replicates are shown for each variant.

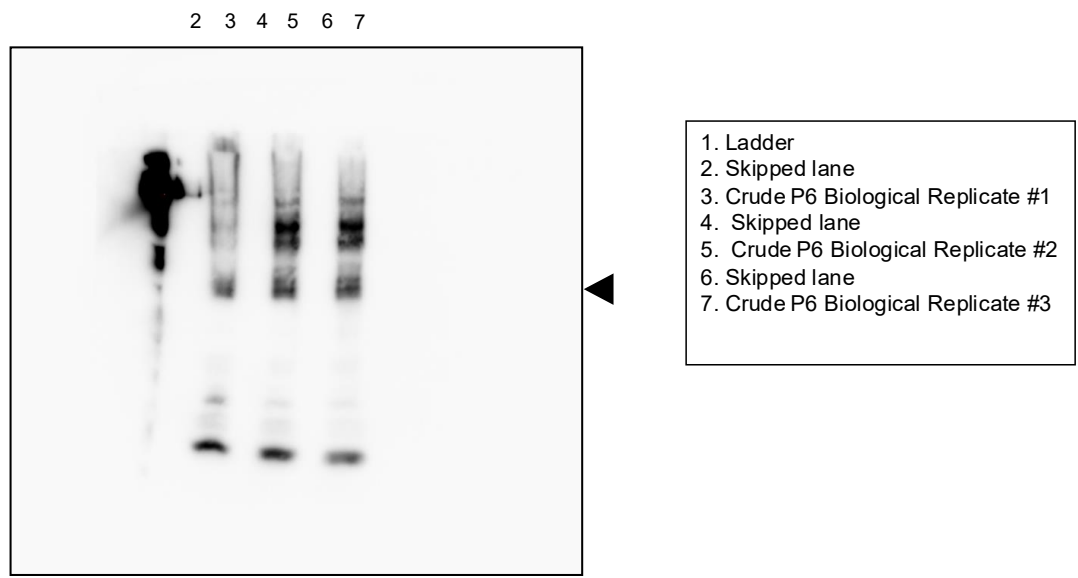

**Fig. S8. Uncropped western blots of patient 6 TMEM165 variants detected by anti-His antibody.** Uncropped western blots showing anti-His detection of the patient 6 TMEM165 variants. Three independent biological replicates are shown for each variant.

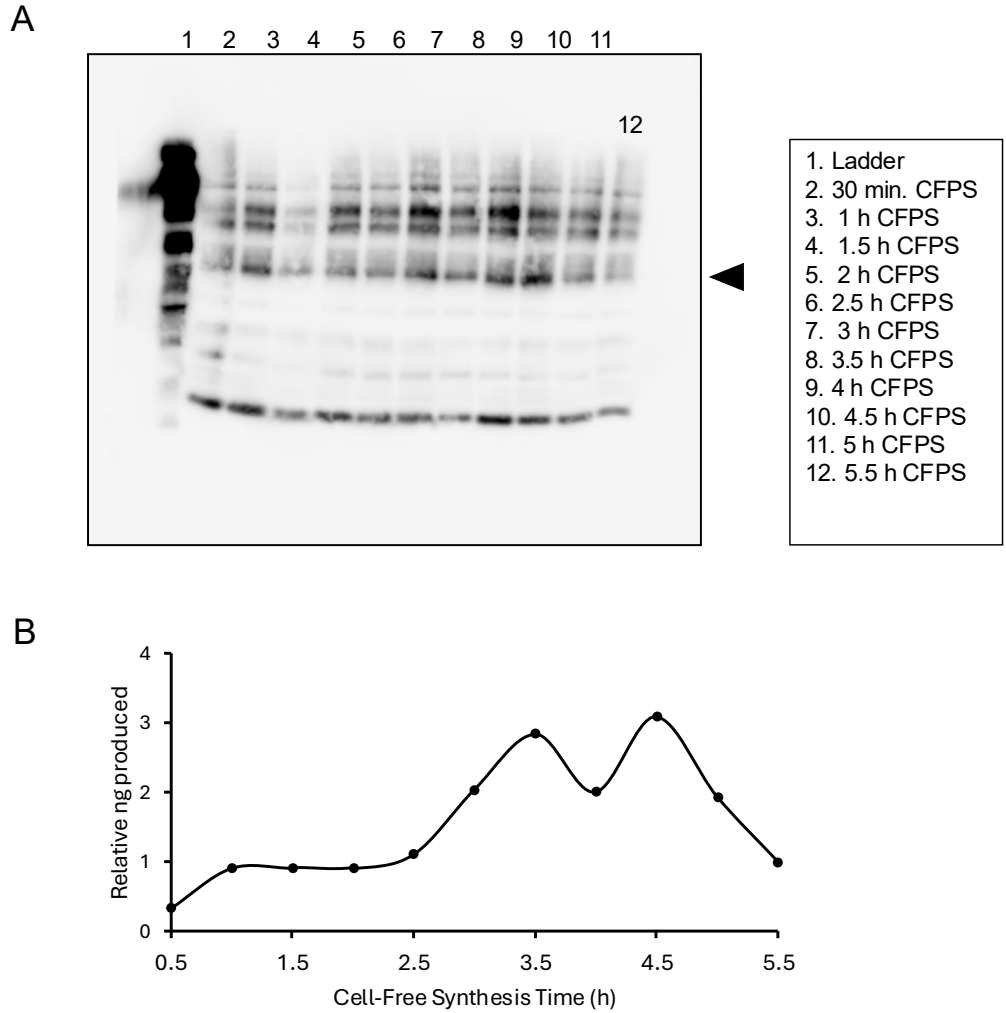

**Fig. S9. Time-dependent synthesis of WT TMEM165.** WT TMEM165 synthesis was assessed over time by western blot using an anti-His antibody. **(A)** Uncropped western blot showing WT TMEM165 expression at each synthesis time point. **(B)** Densitometric analysis of WT TMEM165 synthesis over time. The highest relative yield was observed between 3 and 4.5 h, which was used to guide the synthesis time elected for experiments.

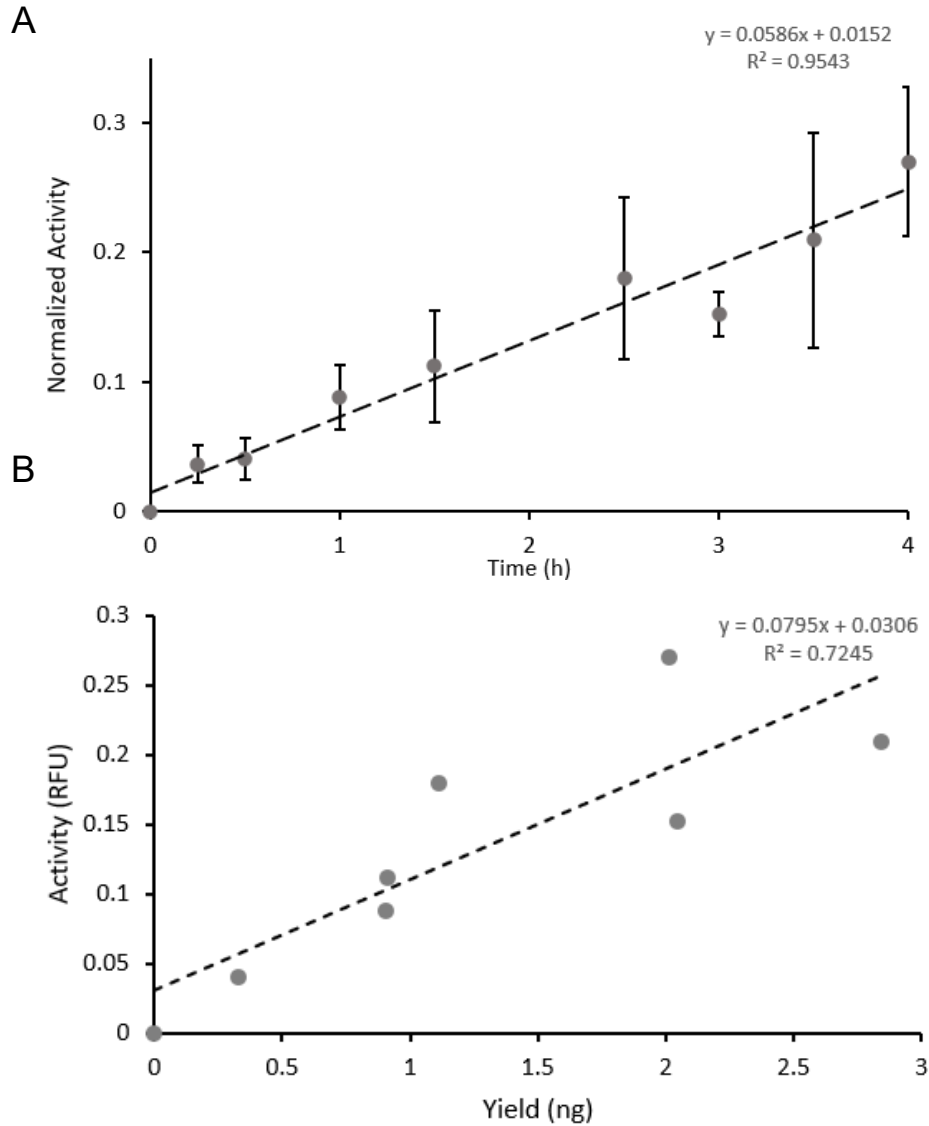

**Fig. S10. Validation of the linear relationship between TMEM165 synthesis yield and transport activity.** Using the synthesis time range identified in Fig. S9, TMEM165 variants were synthesized between 0 and 4 h to evaluate whether measured transport activity scaled with protein yield. **(A)** Transport activity of TMEM165 variants synthesized for the indicated times. Activity was normalized to the Triton X-100 response to account for total encapsulated sensor signal. **(B)** Relative TMEM165 protein yield calculated from densitometric analysis of western blots from the same samples. Both transport activity and protein yield increased linearly over this synthesis range, supporting normalization of mutant transport activity to relative protein expression levels in subsequent analyses.

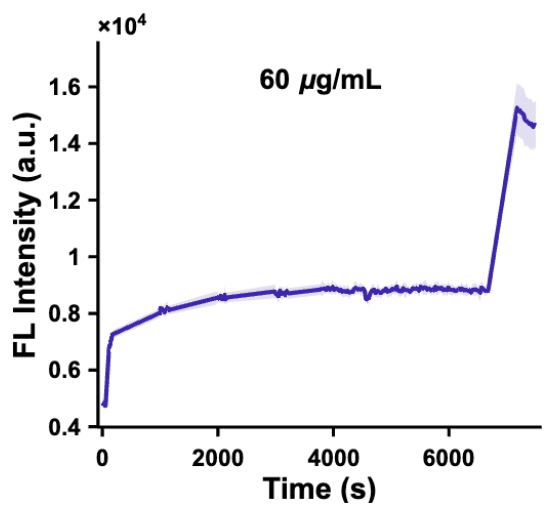

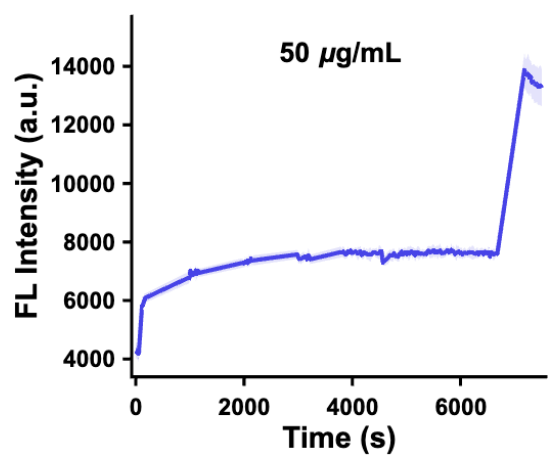

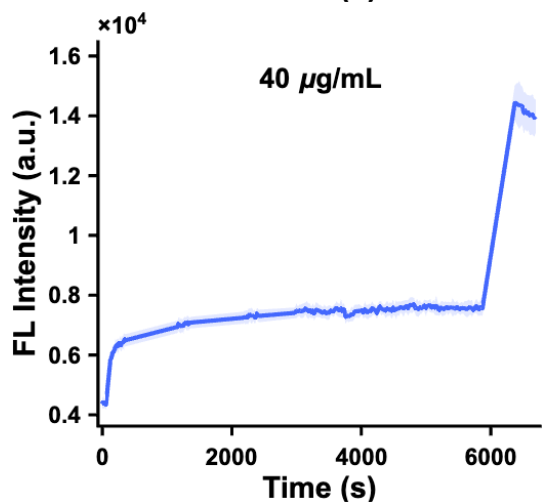

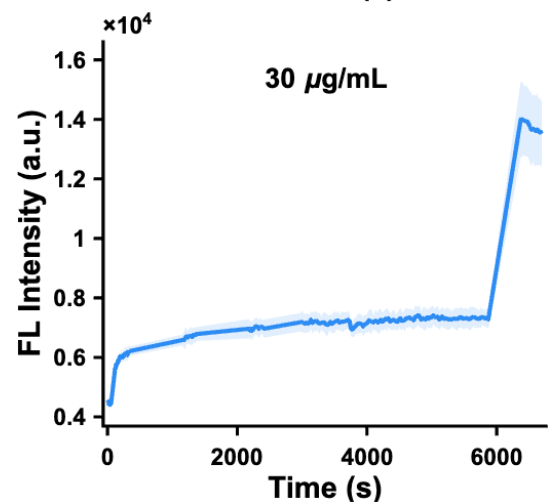

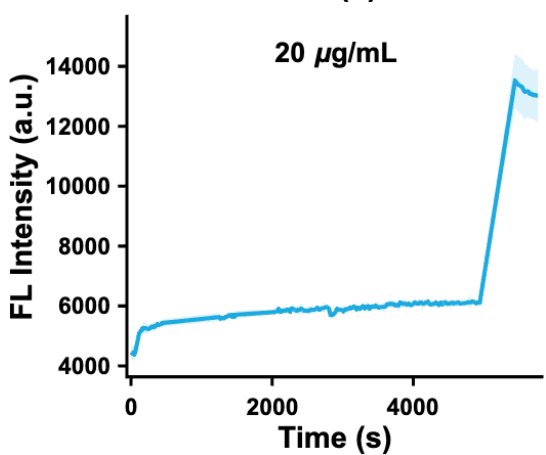

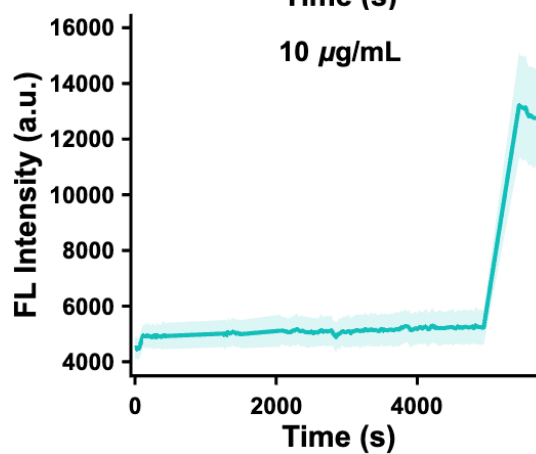

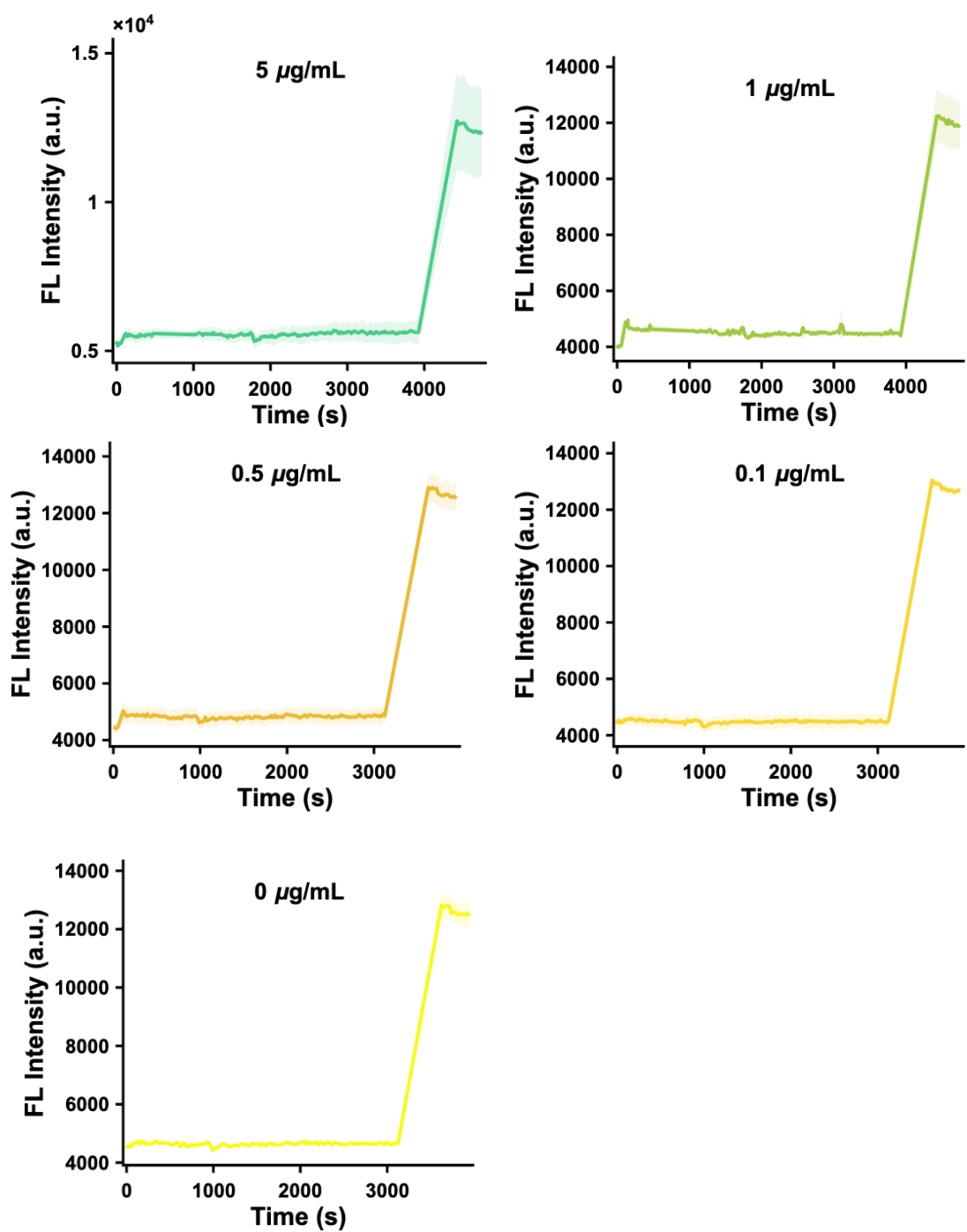

**Fig. S11. Validation of the calcium influx assay using α-hemolysin.** The calcium influx assay described in Fig. 3 was performed using α-hemolysin as a pore-forming positive control. Liposomes containing encapsulated Fluo-4FF were exposed to increasing α-hemolysin concentrations, with 10 concentrations tested across the indicated range. Calcium-dependent fluorescence increased in response to α-hemolysin, consistent with concentration-dependent membrane permeabilization and $\text{Ca}^{2+}$ influx. Three independent replicates are shown for each condition.

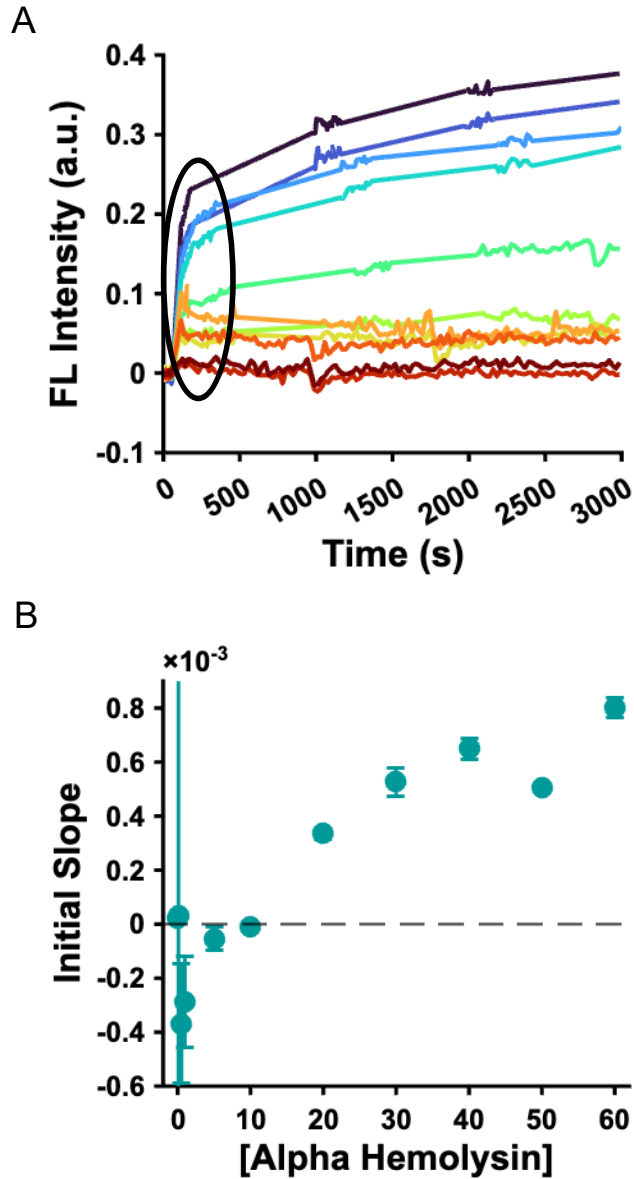

**Fig. S12. Quantitative analysis of α-hemolysin-dependent calcium influx.** Data from Fig. S11 were analyzed to determine the concentration range over which α-hemolysin activity scaled linearly with protein concentration. **(A)** Zoomed-in view of the first 3000 s of the calcium influx assay, prior to Triton X-100 addition. The initial fluorescence jump, indicated by the circled region, is attributed to $\text{Ca}^{2+}$ binding to non-encapsulated sensor; while the subsequent fluorescence increase is attributed to α-hemolysin-mediated $\text{Ca}^{2+}$ influx into liposomes. **(B)** Initial calcium influx signal plotted as a function of α-hemolysin concentration. The lowest concentrations tested, 0, 0.1, and 0.5 $\mu\text{g/mL}$, were noisy and fell below the assay signal-to-noise threshold. The remaining concentrations, from 1 to 60 $\mu\text{g/mL}$, fell within a linear response range. These data define the minimum protein amount required for reliable detection and further demonstrate that measured calcium influx scales linearly with protein amount in an independent pore-forming model system. Error bars represent standard deviation.

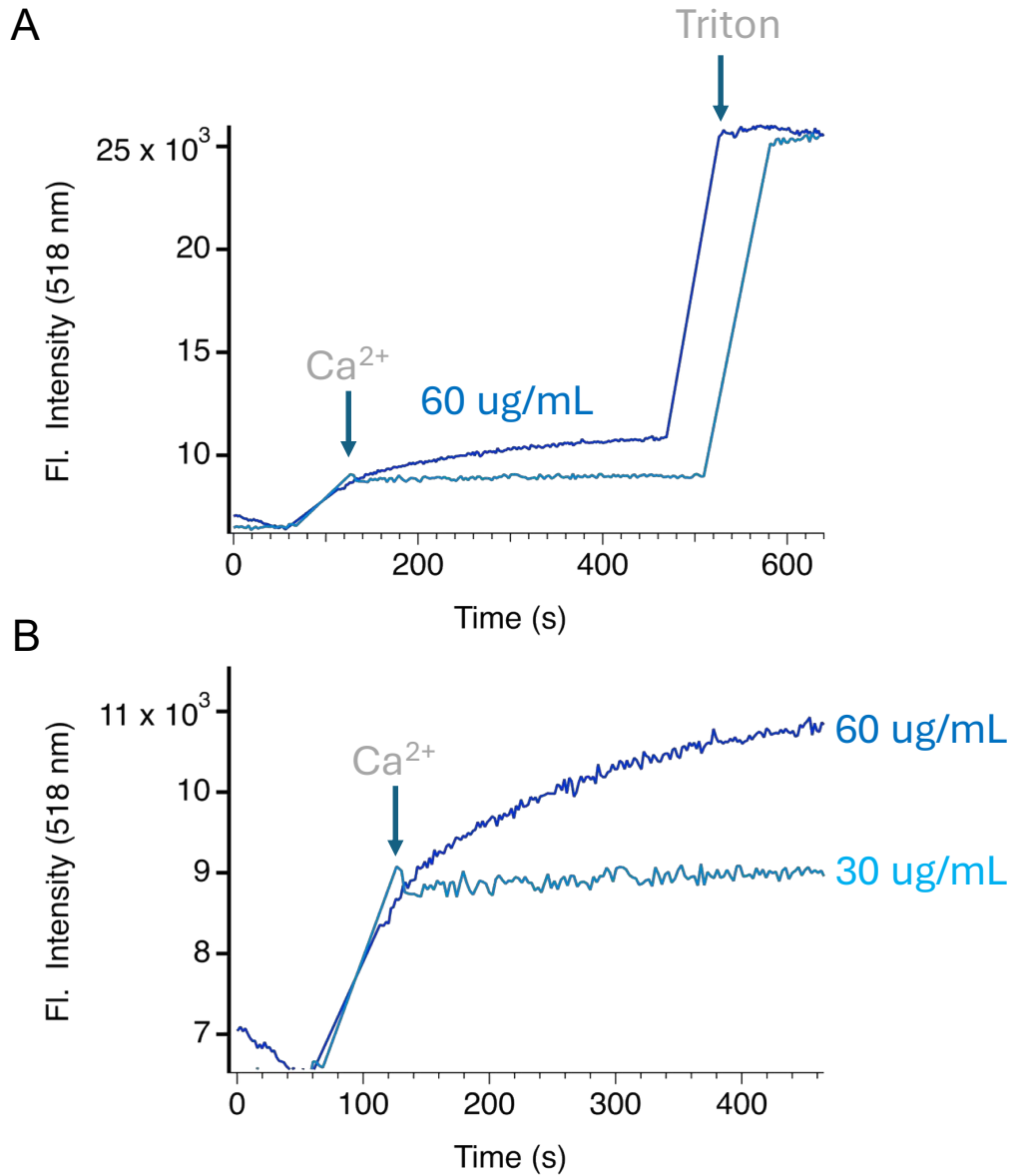

**Fig. S13. Comparison of α-hemolysin-mediated calcium influx at 30 and 60 $\mu\text{g/mL}$.** Calcium influx traces were compared for liposomes treated with 30 or 60 $\mu\text{g/mL}$ α-hemolysin. **(A)** Full fluorescence traces, including Triton X-100 addition, overlaid and normalized to the Triton response. **(B)** Zoomed-in view of the initial fluorescence spike, indicated by the arrow, and the subsequent α-hemolysin-mediated fluorescence increase. The 60 $\mu\text{g/mL}$ condition begins to plateau following the initial influx period, consistent with more rapid $\text{Ca}^{2+}$ equilibration at the higher α-hemolysin concentration.

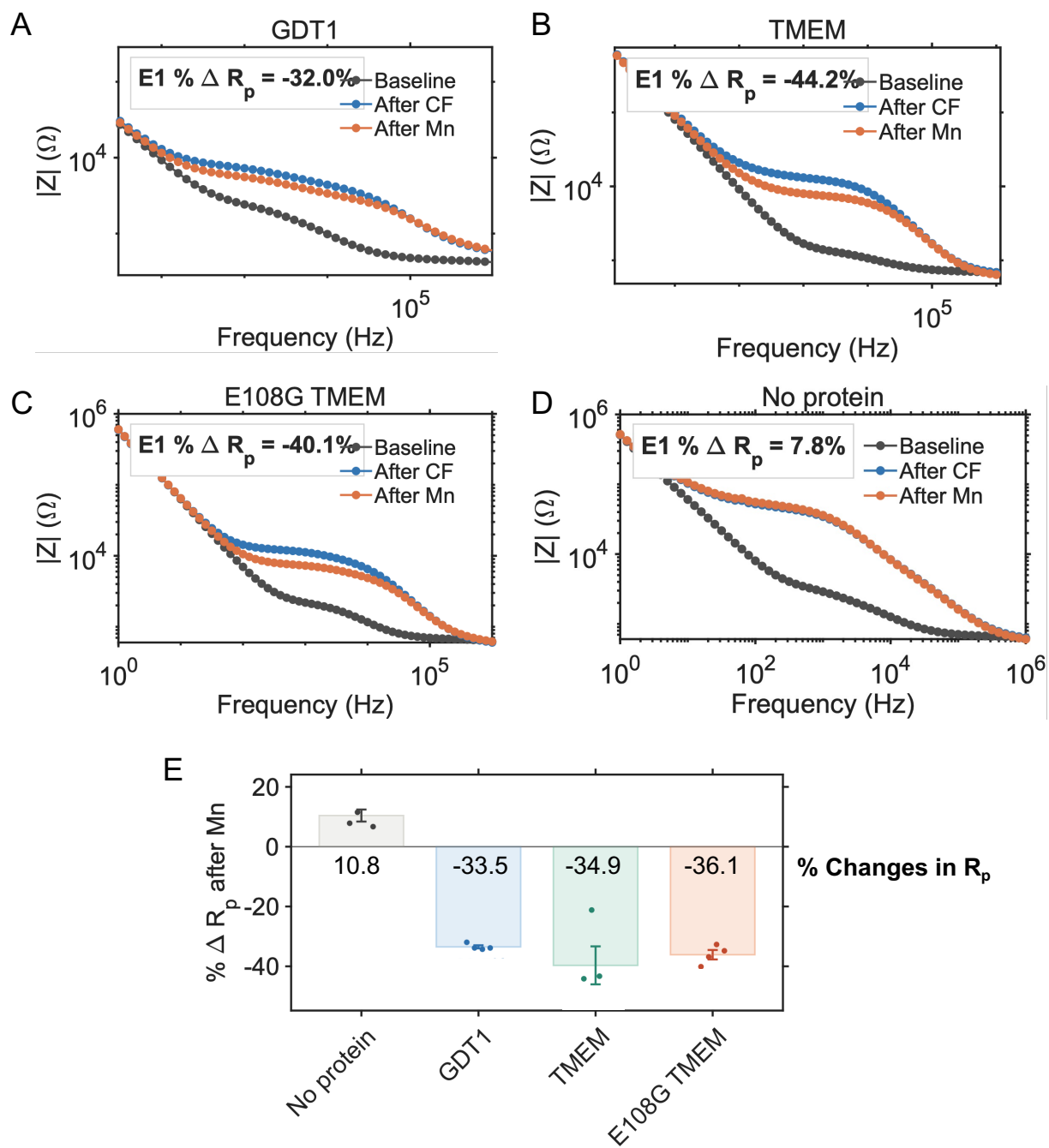

**Figure S14. Mn-dependent changes in EIS response for GDT1, TMEM, E108G TMEM, and no-protein control.** Representative Bode magnitude plots showing impedance magnitude, $|Z|$, as a function of frequency for GDT1 (A), TMEM (B), E108G TMEM (C), and no-protein control (D). For each sample, traces are shown at baseline, after cell-free protein synthesis/assembly (After CF), and after Mn addition (After Mn). The inset value in each plot indicates the percent change in fitted resistance, $R_p$, from After CF to After Mn. (E) Quantification of percent change in $R_p$ after Mn addition. Bars represent the mean percent change, error bars indicate SEM, and individual points correspond to independent electrodes. GDT1, TMEM, and E108G TMEM showed decreases in $R_p$ after Mn addition, whereas the no-protein control showed a small positive change.

A

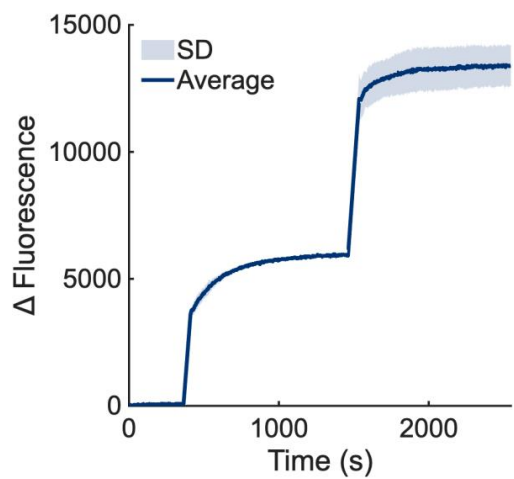

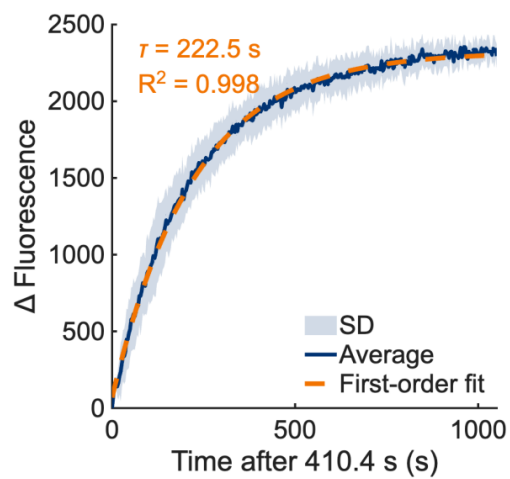

B

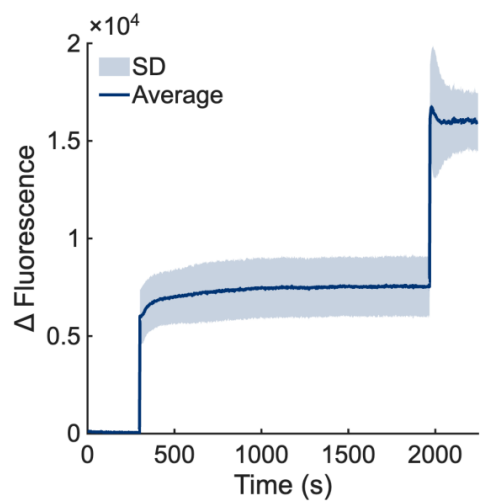

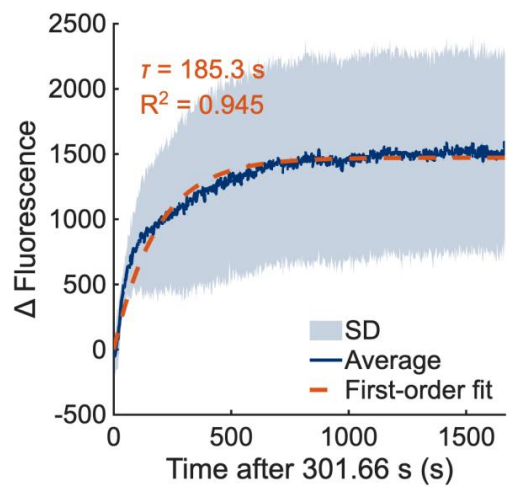

C

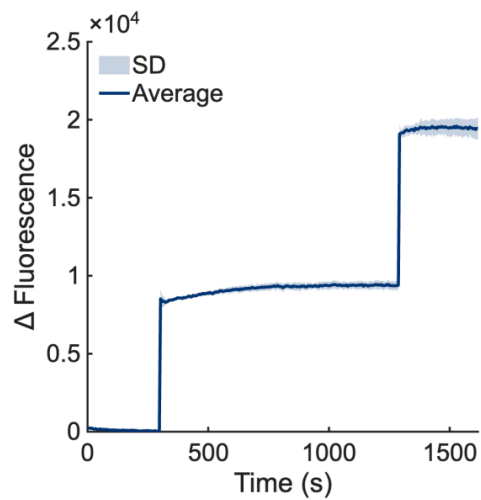

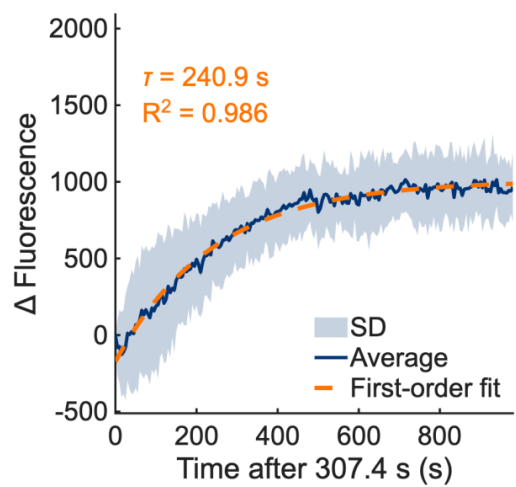

D

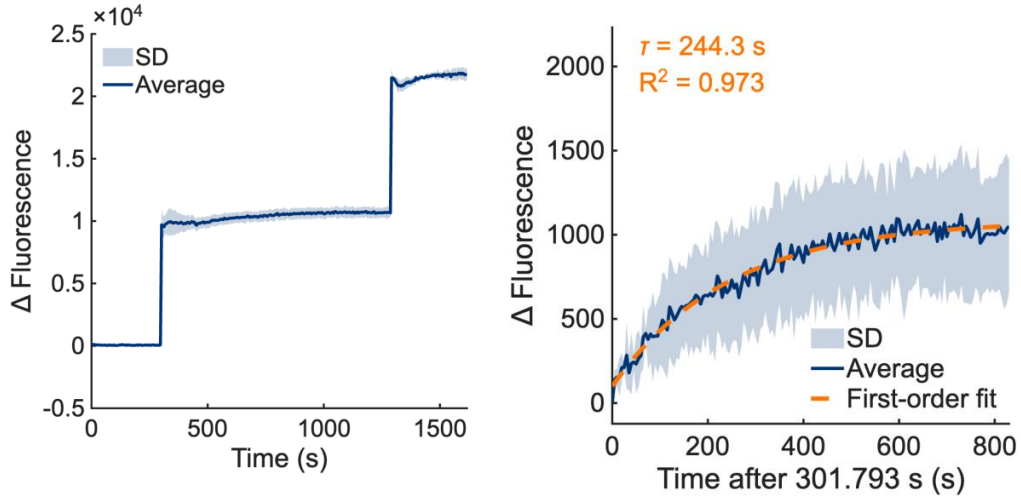

E

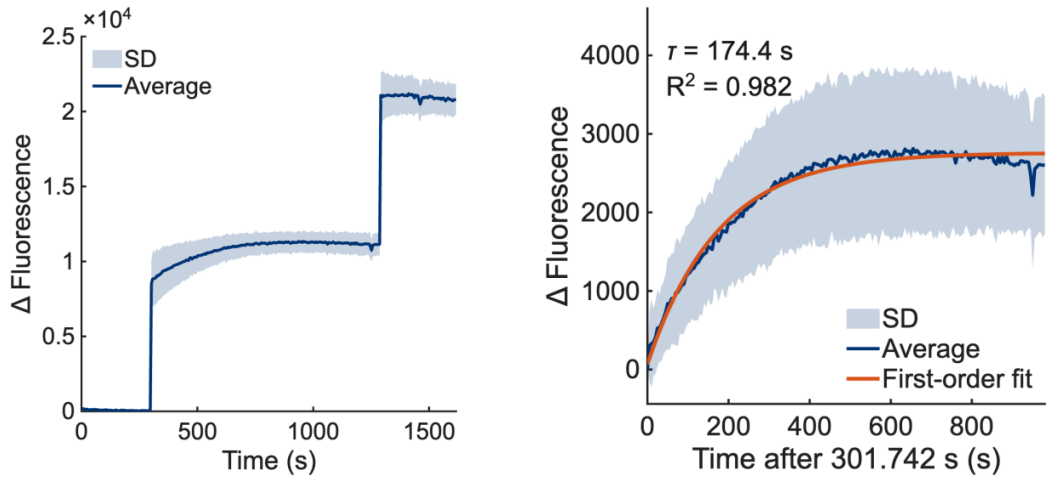

**Fig. S15. Calcium transport activity assays for TMEM165 patient variants.** Calcium influx assays were performed for each TMEM165 patient variant. For each panel, the left trace shows the full fluorescence response, and the right trace shows a zoomed-in view of the flux region beginning at the time point indicated by the arrow in Fig. S13. Solid lines represent the average trace, and shaded regions represent standard deviation across three independent experiments. The zoomed-in flux regions were fit to a single-exponential model, with the fitted rate constant and $R^2$ value displayed on each plot. The experimental average is shown as the solid line, the standard deviation as the shaded region, and the single-exponential fit in orange. (A) P1–P3. (B) P4. (C) P5–304. (D) P5–126. (E) P6.

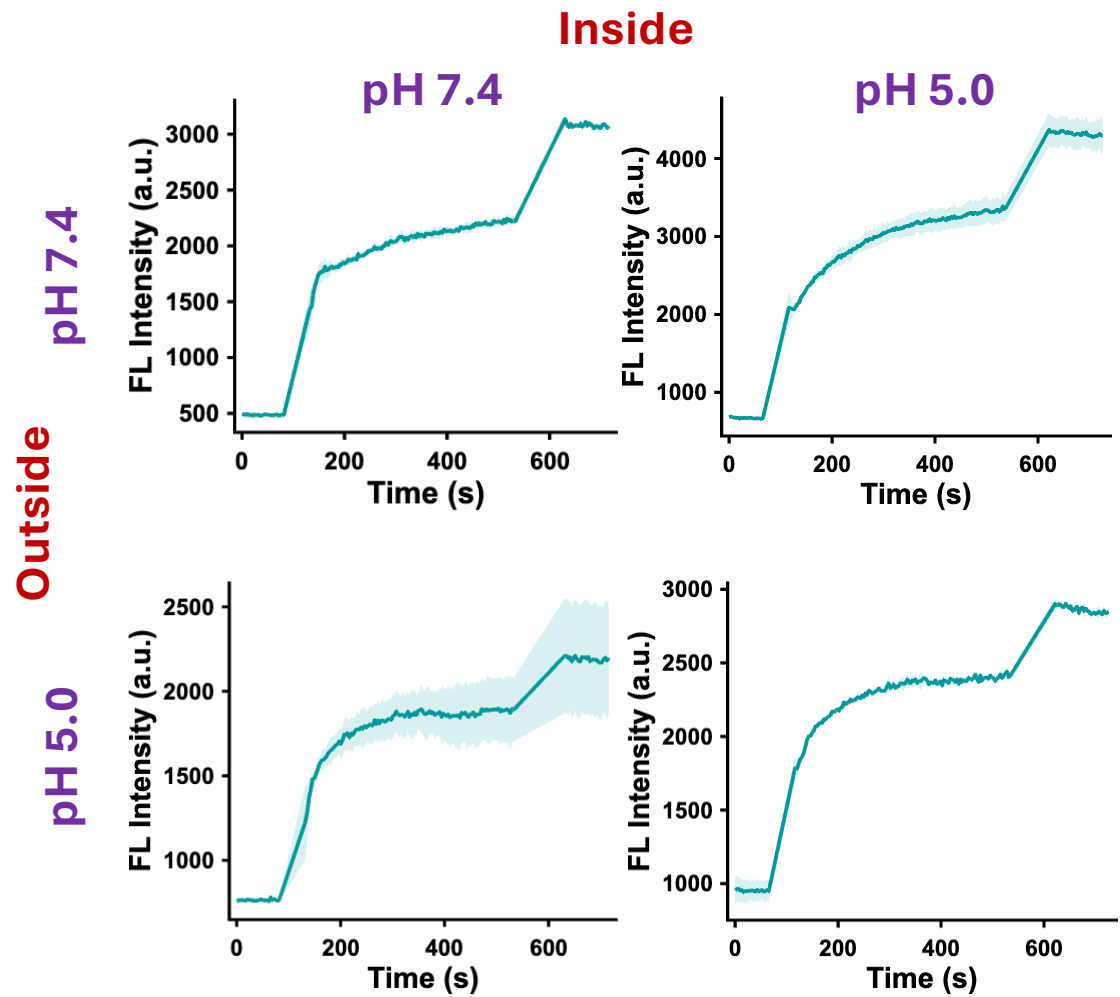

**Fig. S16. pH-dependent calcium transport activity of WT TMEM165.** Calcium influx assays were performed with WT TMEM165 under different internal and external pH conditions. The internal liposome pH is indicated above each column, and the external buffer pH is indicated along the left side of each row. For each condition, three independent full fluorescence traces are overlaid.

**Fig. S17. pH-dependent calcium transport activity of the TMEM165 E108G variant.** Calcium influx assays were performed with the TMEM165 E108G variant under different internal and external pH conditions. The internal liposome pH is indicated above each column, and the external buffer pH is indicated along the left side of each row. For each condition, full fluorescence traces are overlaid.

**Fig. S18. E108G disrupts pH gradient proton transport of TMEM.** (A) FITC fluorescence spectra measured across pH 5-7 demonstrate increasing fluorescence with increasing pH. (B) Proton transport by TMEM 165 was monitored in liposomes with an internal pH of 5 and external of 7. wtTMEM produces an increase in encapsulated fluorescence signal consistent with an increase in internal pH. E108G does not exhibit the same increase suggesting a disrupted transport.

**Fig. S19. Identification of the linear response range of the sodium sensor.** The sodium sensor was calibrated across varying $\text{Na}^+$ concentrations to identify the concentration range suitable for quantitative sodium influx measurements. **(A)** Emission spectra of the sodium sensor collected across the indicated $\text{Na}^+$ concentrations, with three independent replicates overlaid for each condition. **(B)** Relative fluorescence intensity plotted as a function of $\text{Na}^+$ concentration to identify the linear response range of the sensor at 545 nm. Measurements were performed in triplicate, and error bars represent standard deviation.
